# OpenPathSampling: A Python framework for path sampling simulations. II. Building and customizing path ensembles and sample schemes

**DOI:** 10.1101/351510

**Authors:** David W.H. Swenson, Jan-Hendrik Prinz, Frank Noe, John D. Chodera, Peter G. Bolhuis

## Abstract

The OpenPathSampling (OPS) package provides an easy-to-use framework to apply transition path sampling methodologies to complex molecular systems with a minimum of effort. Yet, the extensibility of OPS allows for the exploration of new path sampling algorithms by building on a variety of basic operations. In a companion paper [Swenson et al 2018] we introduced the basic concepts and the structure of the OPS package, and how it can be employed to perform standard transition path sampling and (replica exchange) transition interface sampling. In this paper, we elaborate on two theoretical developments that went into the design of OPS. The first development relates to the construction of path ensembles, the *what* is being sampled. We introduce a novel set-based notation forthepath ensemble, which provides an alternative paradigm for constructing path ensembles, and allows building arbitrarily complex path ensembles from fundamental ones. The second fundamental development is the structure for the customisation of Monte Carlo procedures; *how* path ensembles are being sampled. We describe in detail the OPS objects that implement this approach to customization, the MoveScheme and the PathMover, and provide tools to create and manipulate these objects. We illustrate both the path ensemble building and sampling scheme customization with several examples. OPS thus facilitates both standard path sampling application in complex systems as well as the development of new path sampling methodology, beyond the default.

## I. INTRODUCTION

Many dynamical processes, including nucleated phase transitions, chemical reactions, and complex conformational changes in biomolecular systems, such as proteins and nucleic acids, occuron longtimescales [1–4], primarily due to large kinetic barriers between metastable states [5–7]. Straightforward molecular dynamics simulations are then highly inefficient due to the long waiting times within metastable basins, while the rare events of interest occur over a short time [8]. Methods such as umbrella sampling [9], blue moon sampling [10], local elevation sampling [11], conformational flooding [12], hyperdynamics [13], metadynamics [14], adaptive biasing force methods [15], replica exchange [16], simulated tempering [17], integrated sampling [18], orthogonal space sampling [19], and numerous others enhance the occurrence of the rare event by bi-asingthe potential energy surface orthe density of sampled conformations. To be effective, bias potentials require (a set of) collective variables that approximate the reaction coordinate. However, a poor choice will lead to poor sampling of reactive pathways and hence poor estimates of the dynamical bottlenecks and the related barrier heights and rates.

The transition pathsampling(TPS) methodology [20–23]) can enhance the sampling of complex dynamical transitions in complex (bio)molecular systems, avoiding the exponentially long timescales that the system spends in metastable states, and, most importantly, bypassing the need fora reaction coordinate. Notwithstanding the efficiency of path sampling, the complexity of implementation and lack of standard tools have hampered widespread application.

The OpenPathSampling (OPS) framework aims at providing a toolbox to make complex transition path sampling simulation easy accessible for users. In Paper I, we introduced the basic concepts and structure of the OPS framework, discussed its ingredients and gave a tutorial on how to conduct some standard simulations using the OPS framework [24]. The current work builds heavily on this companion paperand we referthe readerto Ref. [24] for full details. The OPS framework facilitates implementation of the three stages in any path sampling study: initialization, sampling, and analysis. In the initialization step the user defines the (network of) transitions that needs to be sampled. This requires definition of stable states and path and/or interface ensembles to be used in (replica exchange) transition interface sampling (TIS) schemes. These definitions are based on phase space volumes defined as a function of a priori chosen collective variables. Since path sampling is basically a Monte Carlo (MC) approach, the user also has to decide on the specific details of how each type of move is implemented, which OPS facilitates with the `MoveStrategy` objects, and the user has to create the overall (and sometimes complex) decision treeforthe MC procedure, which OPS implements in a `MoveScheme`. The transition network and the `MoveScheme`, together with the Molecular Dynamics Engine, a Storage file, and a initial path sample set, enterare used by the `PathSimulator`, which performs the sampling. Analysis of the sampled paths is subsequently performed using information obtained from the Storage object. OPS provides tools for the initialization step and for the analysis. For more details we refer to Paper I [24] and to the online documentation at http://openpathsampling.org.

This paper is mainly aimed at method developers and researchers interested in devising their own path sampling methods using the OPS framework. This requires an extensive treatment of the more fundamental ideas that went into the design of OPS. The paper focuses on two of those fundamental aspects. One is the construction of path ensembles, which can be viewed as *what* is being sampled. The other is the customisation of the Monte Carlo procedures, which relates to *how* the path ensembles are being sampled. This paper provides novel conceptional frameworks for dealing with these two aspects.

In the first part, we focus on the path ensembles. While the definition of path ensembles in the original TPS and TIS papers is perfectly usable for many applications, these definitions can be come quite cumbersome when multiple states or multiple collective variables come in to play. This also holds for the more complex path moves, such as the *minus interface move* [25, 26] used in replica exchange TIS (RETIS), which exchanges a trajectory in the first interface ensemble with a trajectory exploring the stable state (the minus interface ensemble) in order to decorrelate the (usually short) pathways in the first interface, and to provide a direct estimate for the flux out of the stable state [25–27]. Here, we present a framework allowing one to build arbitrarily complex path ensembles from fundamental path ensembles. To facilitate this, we first introduce a novel set-based notation for the path ensembles. For completeness, we also provide connections to the original TPS and TIS notation. This novel notation provides an alternative paradigm for constructing path ensembles with several major advantages: (1) It allows one to easily create complex ensembles as combinations of simpler ensembles, e.g., using set logic. (2) It creates a systematic connection betweentheensemble indicatorfunction and the stopping criteria used when generating a trajectory for the ensemble, e.g., with a shooting move. Previously, the stopping criteria were identified separately for every ensemble/path generating move. (3) It facilitates analysis, as many analysis procedures can be framed as searching for subtrajectories that satisfy some ensemble indicator function. Examples of this are provided in Sec. IV. Of particular importance herein is the sequential path ensemble, which is directly related to the way that OPS implements path sampling and testing. We explain in detail how different ensembles are being built in terms of these sequential path ensembles. We end the first part with a set of general guidelines and simple rules on how arbitrary path ensembles could be built in OPS.

In the second part of the paper, we describe the framework that creates the Monte Carlo process used by OPS. This framework is designed to be extremely flexible, which enables one to customize the Monte Carlo move scheme and to build non-standard path sampling schemes with little effort. This ability to customize the move scheme is one of the major advantages of the OPS framework. It allows experiences users to design a sampling method tailored to a specific system. Here the two central concepts are (1) the move scheme, which encodes the entire Monte Carlo procedure as a decision tree, and (2) the path movers, which perform the moves. We describe both concepts in detail, as well as the tools in OPS that facilitate customization of the move scheme.

The paper is organized as follows. In Sec. II A we briefly review the original standard notation for TPS path ensembles. Subsequently, we introduce the novel path ensemble set notation, including the sequential ensemble. We then describe in Sec. II E how OPS implements these ensembles, and give some guidelines and rules on how new ensembles could be built. In Sec. III we discuss customizing the Monte Carlo moves in detail. We give illustrations of these concepts in Sec. IV, where we discuss the application of generating and splitting trajectories, as well as customizing the move schemes for alternative replica exchange simulations. Finally, we end with conclusions and an outlook.

## II. BUILDING BLOCKS FOR PATH ENSEMBLES AND VOLUMES

### A. Standard TPS and TIS notation

In this section, we briefly recapitulate the standard notation for the path ensemble and distribution functions used in TPS and TIS before introducing the novel set based notation that is more commensurate with the way OPS implements path sampling. This section is not meant as an introduction to path sampling, but rather to describe the connection between the novel set-based notation and the standard notation found in the literature. For a review of path sampling methodology we refer the reader to Refs. [20–23, 28]. In the next sections we follow the notation that was introduced in Refs. [26] and [28].

#### 1. The TPS path ensemble

A path is a discretized, time-ordered sequence of states in phase space x ≡ {*x*_0_, *x*_1_; *x*_2_,…, *x*_L_}, in which consecutive states, or frames, are separated by a small time increment Δ*τ*. Each frame *x* = {*r, p*} consists of the positions and momenta of all particles in the entire system. The path-length *L* can be chosen fixed or variable, depending on the type of path ensemble. The path probability for a trajectory of duration 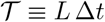 is

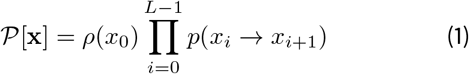

where *p*(*x*_*i*_ → *x*_*i*+1_) denotes the Markovian transition probability to go from *x*_*i*_ to *x*_*i*+1_ in one time step, which depends on the underlying dynamics [22]. Further, *ρ*(*x*_0_) is the distribution of initial conditions, in many cases the equilibrium distribution. TPS constrains pathways between two stable states A and B. Such states are defined using an collective variable or order parameter *λ*, for example [^1^ Note that the term ‘order parameter’ and ‘collective variable’ are used interchangeably in this work. As explained in Paper 1 [24], the term collective variable refers to any function of the system’s coordinates.]

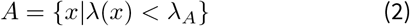

and likewise for B. Here, *λ*(*x*) returns the order parameter λ for frame *x*, and *λ*_*A*_ defines the boundary of state A.

The standard TPS path ensemble distribution with a fixed length *L* constrains the path to begin in *A*, and end in *B*

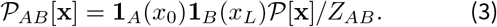

where 1_*A*_(*x*_0_) and 1_*B*_(*x*_*L*_) are indicator functions that are, respectively, unity if the trajectory starts with *x*_0_ ∈ *A* and ends with *x*_*L*_ ∈ *B* and zero otherwise. The formal definition of 1_*A*_(*x*_0_) is

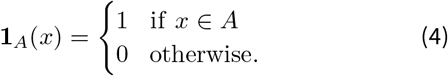

and 1_*B*_(*x*_*L*_) is defined likewise. The normalization factor 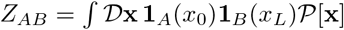 is akin to a partition function, where the integral over 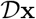 takes into account paths of length L starting at all possible initial conditions *x*_0_.

For variable path length TPS a similar path ensemble distribution can be written

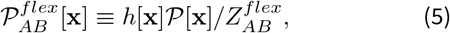

where the indicator function *h*[x] now selects the paths that immediately leave *A*, and just enter *B*

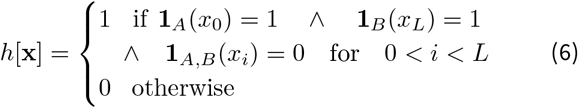

Note that this path ensemble indicator function already shows some complexity. The normalization factor is now 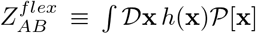 where the integral over 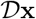 takes now into account paths of all length starting at all possible initial conditions *x*_0_.

#### 2. The TIS path ensemble

Transition Interface sampling (TIS) defines a series of successive non-intersecting interfaces *λ*_0_, *λ*_1_,…, *λ*_*n*_, based on an order parameter *λ*, and samples the TIS path ensemble for each interface. Paths in the interface ensemble *i* start in A (at *λ*_0_), cross the interface *λ*_*i*_ at least once, and finally either return to A, or end in B. Defining adjacent phase space regions separated by interface *λ*_*i*_ as 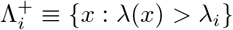,the path probability for an interface ensemble *i* is given by

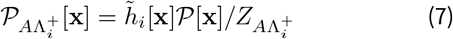

where the subscripts 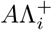 denote the phase phase regions connected by the paths, and the TIS indicator function

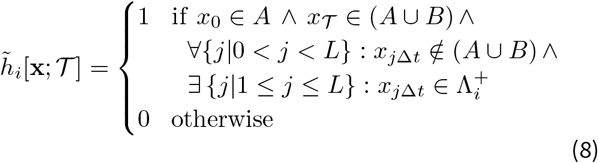

where the first and second line ensures that only the initial and end pointsare in Aand B, respectively, whereasthethird line requires that the path cross the interface. The normalization factor 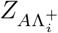 is defined by

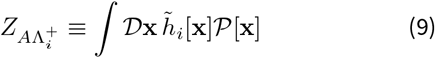

where the path integral runs again overall possible paths of all lengths. The ensembles for the reverse reaction *B* → *A* are defined in an analogous fashion [26, 28].

The path ensemble indicator functions become rather complex even for the basic TPS and TIS ensembles. Moreover, the indicator function as described above are not directly implementable in a path sampling code such as OPS, as they only apply to entire paths. During a shooting move, which is at the heart of TPS and TIS, OPS has to monitor a newly generated path and apply a stopping criterion. Such a criterion requires a notation that is better suited to the way that OPS implements both the monitorfunction and the path ensembles themselves.

### B. Volumes as sets

As the novel notation is based on set logic, it is only natural to also treat the stable states as sets. In OPS, stable states, and in fact any region in phase space, are described as “volumes”. Additionally, TIS interfaces are treated as volumes. This has several advantages, among which is the fact that it is then easy to combine volumes using set logic.

Volumes are defined by collective variable functions. As mentioned above, a state consists of the (infinite) set of all configurations that obey the state definition. For instance, using one order parameter *λ*(*x*) state A can be defined as

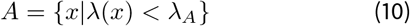

More general definitions are possible, e.g. by using an arbitrary number of order parameters. TIS interface volumes for an interface *i* connected to state *A* can be defined as

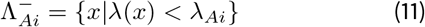

which is the part of phase space complementary to 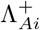 defined in the previous section. Crossing an interface now amounts to leaving 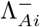, putting it conceptually on the same level as leaving A or entering B.

Volumes (e.g., states) can be combined using set logic. For two states A and B

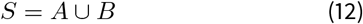

denotes the union of sets, while

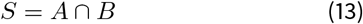

denotes the intersection. In this way volumes obeying an arbitrary number of conditions can be constructed.

In OPS this set logic is implemented by several functions. We can take unions and intersections of volumes, and negations of volumes using the objects `UnionVolume`, `InteractionVolume` and `NegatedVolume`. From these operations any logical operation can be constructed. Take as an example the `SymmetricDifference`, which fortwo volumes A and B would amount to all points that are either in A or in but not in both. This is logically equivalent to

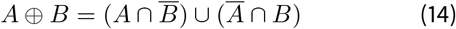

where the overbar denotes the complement or negation of the volume. Other logical operations can be constructed at will.

### C. Path ensembles as sets

#### 1. Unifying two basic tasks in OPS

As discussed in Sec. II A, path ensembles are the set of all trajectories that satisfy the ensemble indicator function *h*_*E*_ [x] of the ensemble *E*, weighted by the natural path probability 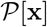. In OPS, the ensemble indicator function is part of the Ensemble object. Paths are sampled with the correct relative weights by using Monte Carlo moves that preserve the distribution. Trajectories with non-zero weight form a set of paths that satisfy the constraints imposed by the ensemble indicatorfunction. Indeed, these constraints define the ensemble indicatorfunction. Forinstance, a TPS ensemble requires the constraints that the first snapshot be in the initial state, that the last snapshot be in the final state, and that no other frames of the trajectory visit a stable state. An important task in OPS is to test whether a trajectory fits a certain ensemble. The Ensemble object takes a trajectory as input and returns whether or not it belongs to that ensemble. Take, for instance, the simple ensemble for which all frames should be in a certain state *A*. The predefined ensemble class `AllInXEnsemble(state)` tests exactly that, returning True only if all frames are in the given state. One of the most productive ways to define useful path ensembles in OPS is the `SequentialEnsemble` object. `SequentialEnsemble` comprise a list of path ensembles that the trajectory needs to fulfill in the correct order. This is crucial when performing path sampling, identifying whether a path fulfills the right conditions for a move, e.g. an exchange move. Moreover, they can be useful foranalysis of pathways.

When performing path sampling, and in particular during a shooting move, another important task in OPS is to monitor whether a trajectory is finished, i.e., fulfils the conditions forstopping. Atfirstsight, one mightthinkthatsuch a test is simply applying the same `Ensemble` object as above. However, this is not the case, since there are obviously many paths that do not obey the desired path ensemble, but still are clearly to be rejected. For instance a path that leaves *A* and returns to *A*, without having visited *B*, is clearly to be rejected. Hence, a halting criterion is needed, or rather, a continuing criterion that tells OPS to keep integrating the molecular dynamics trajectory, until it is clear that the trajectory can no longer ever satisfy the path ensemble. The can_append method provides the test for whether the path can be extended or not. Moreover, the can_append method is the building blockfrom which the `SequentialEnsemble` is constructed.

In the following sections, we present a general set-based approach, which connects the ensemble to its halting criteria, and which allows one to build arbitrarily complex ensembles from simple building blocks. As we build up to more complicated ensembles, we will, at each stage, first describe the set-based approach, introducing a new notation for describing path ensembles. Then we will show how that new notation maps directly onto objects in OPS.

#### 2. The basic building blocks

As above we denote a (sub)trajectory as a discretized time-ordered sequence of phase space points x ≡ {*x*_*b*_, *x*_*b*+1_,,…, *x*_*e*_} where *b* and *e* denote the beginning and end of the (sub)trajectory, respectively. For *e* < *b* we define the trajectory of zero length x = {}. Note that a time-reversed path also has positive time-order {*y*_0_, *y*_1_;…, *y*_*L*_}, and can be constructed from the trajectory {*x*_0_, *x*_1_,,…, *x*_*L*_} by setting *y*_*i*_ = *x*_*L*−*i*_.

A path ensemble is an (infinite) set of trajectories obeying a certain criterion, encoded by indicator functions. A basic example is the ensemble of trajectories for which all frames are within volume A. In OPS, indicator functions determine whether a trajectory belongs a particular ensemble. For instance, the (formal) function In_*A*_(x) returns unity only when the entire trajectory belongs to volume *A*,

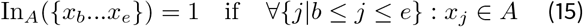

and zero otherwise. Likewise the indicatorfortheset of trajectories entirely outside of *A* requires an indicatorfunction Out_*A*_(x) that determines that no element belongs to *A*

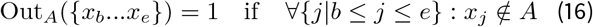

As can be seen directly from these definitions,

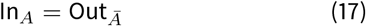

Just as volumes can can be seen as sets allowing set-based logic, path ensembles can becombined orintersected using set logic. An ensemble can be combined, e.g. using a union (indicated by ∪) or an intersection (indicated by ∩). A union of ensembles means that the trajectory has to belong any one of the ensembles; an intersection means that the trajectory has to belong to all ensembles. Combination of these logical operations are likewise defined.

Suppose that we are interested in the ensemble Out_*S*_. with *S* = *A* ∪ *B* the union of states *A* and *B*. The ensemble logic gives

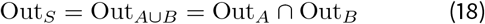

Note that here, the notation In_*A*_ and Out_*A*_ refers to the ensemble, i.e., the entire set of trajectories, whereas when we talk about the associated indicator function we use In_*A*_(x) and Out_*A*_(x).

To construct all possible logical statements, we need the complement or (negation) of ensembles. We can take complements of ensembles, e.g.,

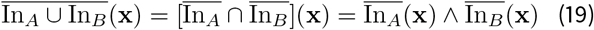

where the overbar indicates the complement of the set. The 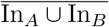 refers to the complement of the union of the se of trajectories entirely inside *A* and the set of trajectories entirely inside *B*. Indeed, a trajectory that is not entirely inside *A* or entirely inside *B*, has to be partly outside *A* and parth outside *B*.

The complement of the Out_*A*_ ensemble is the PartIn_*A*_ en semble, defined by the indicatorfunction:

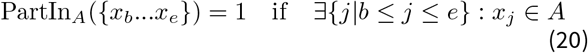

Likewise, the complement of the In_*A*_ ensemble, called PartOut_*A*_, is defined by the indicator function where part of the trajectory is outside *A*.

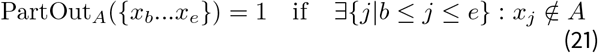

From these definitions it is clearthat

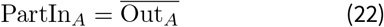

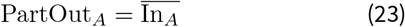

In addition, analogously to Eq. (17), we have

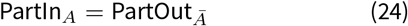

One might expect that In_*S*_ would obey logic analogous to Out_S_. However, it turns out

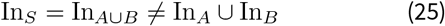

because this would state that either all frames are in *A* or all frames are in *B*. Instead, it is possible that some frames are in *A* and some frames are in *B*, but no frames are outside of *S*. Thus the connection is

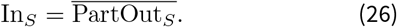

In OPS the four basic functions In_*X*_, Out_*X*_, PartIn_*x*_, PartOut_*X*_ (illustrated in Fig. 1) have their own predefined objects `AllInXEnsemble`, `AllOutXEnsemble`, `PartInXEnsemble`, `PartOutXEnsemble`, which act as building blocks from which ensembles can be constructed. Indeed, as the names suggest, these ensembles only return True if, respectively all frames are in X, all frames are out of X, at least one frame is in X, at least one frame is out of X (see Table I).

**FIG. 1.**
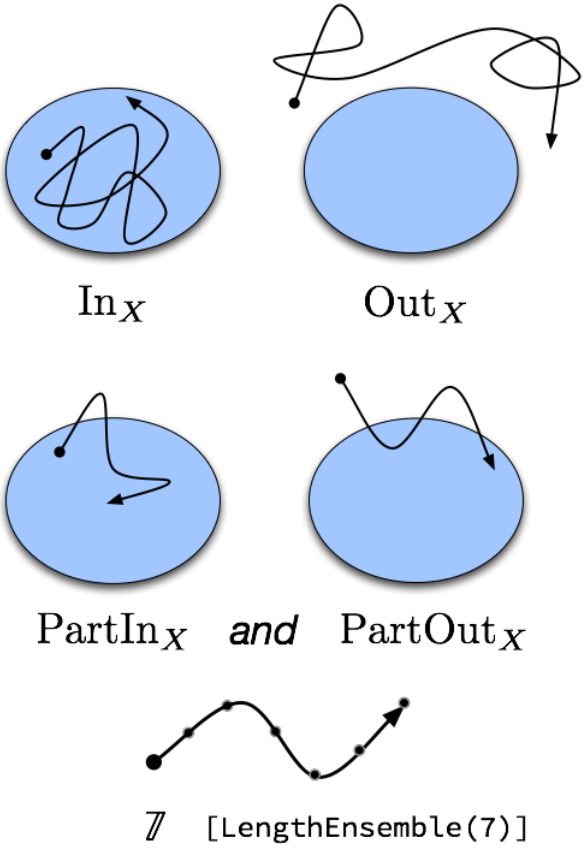
**“Building block” ensembles in OPS**, with example trajectories. Sequences of logical combinations of these ensembles are used to create path ensembles used in OPS. Note that several example trajectories could satisfy the PartIn_*X*_ and PartOut_*X*_ ensembles. Either of the trajectories shown for one would satisfy the other. In addition, the trajectory for the In_*X*_ would also satisfy PartIn_*X*_, and the trajectory for Out_*X*_ would also satisfy PartOut_*X*_.

**TABLE I.** Basic building block ensembles. For each ensemble, the table gives themathematical notation used in Sec. II, the associated OPS class name, a description of the constraint it represents, and the logical complement ensemble.

| Notation | OPS object | Returns True if | Complement |
| --- | --- | --- | --- |
| $\text{In}_X$ | AllInXEnsemble | all frames are in volume $X$ | PartOut$_X$ |
| $\text{Out}_X$ | AllOutXEnsemble | all frames are in volume $\bar{X}$ | PartIn$_X$ |
| $\text{PartIn}_X$ | PartInXEnsemble | at least one frame is in volume $X$ | Out$_X$ |
| $\text{PartOut}_X$ | PartOutXEnsemble | at least one frame is in volume $\bar{X}$ | In$_X$ |
| $\mathfrak{n}$ | LengthEnsemble | trajectory consists of $n$ frames | Any length but $n$ |
| $[\mathbb{E}_1 \rightarrow \dots \rightarrow \mathbb{E}_n]$ | SequentialEnsemble | subtrajectories are in subensembles $\mathbb{E}_1, \dots, \mathbb{E}_n$ in the correct order | Failing any subensemble |

While `PartInXEnsemble` and `PartOutXEnsemble` at first sight seem to be identical ensembles, they are in fact different since `PartInXEnsemble` also yields True for a trajectory that is all in X, whereas `PartOutXEnsemble` does not. Also, contrary to what one might naively think, the complement of `AllInXEnsemble` is not the `AllOutXEnsemble`. As discussed above, the complement of the In_X_ implementation `AllInXEnsemble` is the PartOut_X_ implementation `PartOutXEnsemble`. Indeed, `PartOutXEnsemble` gives True always if one frame is out of X., and only returns False if all frames are in X, the very definition of `AllInXEnsemble`. In Table I the complements of the basic building blockensem-bles are given.

#### 3. The length ensemble

The length ensemble consists of all paths of a specific length *n*. Formally, it can be defined by the indicator function

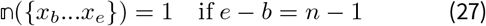

where the 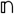 symbol can take any positive integer number *n* > 0. An additional definition is needed forthe zero length *n* = 0 ensemble

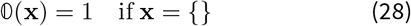

which is the case for *e* < *b*. The indicator function 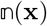 thus returns unity only if the trajectory consists of *n* frames. To test whether a trajectory is entirely in state *A* with a length *n* = 7 thus becomes

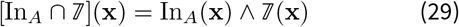

OPS implements this ensemble with the `LengthEnsemble(n)` which, as one might expect, requires the (sub) trajectory to be of a specified length *n*.

### D. The can-append criterion

In practice, path sampling uses methods like the shooting move to generate new trajectories by running dynamics. The shooting move must have some criterion to determine when to stop the trajectories it generates. In early versions of path sampling, this was based only on trajectory length, but more advanced variants gain efficiency by stopping the simulation based on information from the coordinates/momenta of snapshots in the trajectory; for example, stopping upon entering a stable state. As such, each path ensemble must be associated with a halting criterion.

In the formalism presented here, the halting condition (or more correctly, the not-yet-halt condition) is called the can-append criterion. The can-append criterion is associated with and determined by a specific ensemble 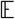 and denoted 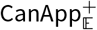. It takes a trajectory x as input and answers the question, “Is the trajectory x a subtrajectory of any trajectory 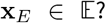” More formally, it is defined as an indicator function on the set of all trajectories for which an additional slice in the forward time direction would not fail the specified criterion for ensemble 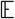. For each ensemble object in OPS, there is a method called can_append that returns True for trajectories that satisfy the can-append criterion, and False fortrajectoriesthatdo not. Forthe negative time direction, there is an analogous criterion 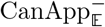 and a can_prepend method, which tests the addition of an extra frame at the beginning of the trajectory. The discussion that follows refers to the forward-time can_append method, but also applies to the backward-time can_prepend.

The indicator function for these ensembles act on a (sub)trajectory x. Perhaps the simplest example is the can-append criterion fora length ensemble:

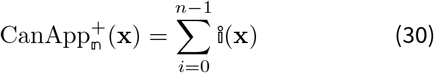

As long as the length of the trajectory is less that *n*, the can-append criterion is satisfied, and `LengthEnsemble.can_append` returns True.

Another example is the In_*A*_ ensemble. The indicator function for 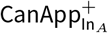 is given by

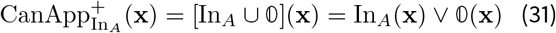

because adding an additional frame that is not in *A* will immediately fail the ensemble. The additional logical or with the zero-length trajectory signifies that for an empty trajectory CanApp should return True, as adding a frame to an empty trajectory is always possible. An analogous formula can be written for 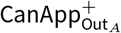.

In contrast, for the ensembles 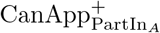 and 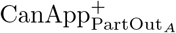, the indicator functions always return True since there is no reason to stop if the additional frame is not in (or out) the state. If a frame of the trajectory is already outside the volume, there is still no reason to stop the trajectory: all extensions will then lead to trajectories that still satisfy the ensemble.

More complex can-append criteria can be constructed using set logic involving intersections and unions [29], or using sequential ensembles, as described below. One important, but somewhat subtle, point is that the logical negation of the result of can_append for an ensemble *E* is not necessarily equal to the result of can_append forthe complement of ensemble *E*. For example, the can_append method for `AllInXEnsemble` returns True if and only if all frames of the input trajectory are in the volume associated with the ensemble. However, as discussed above, the complement of an `AllInXEnsemble` is a `PartOutXEnsemble`, for which the can_append method always returns True. Taking the complement applies to the ensemble; determining the result of the can_append for the complement depends on the complement ensemble, not on the result of can_append in the original ensemble.

OPS also implements two other related methods for each ensemble: `strict_can_append` and `strict_can_prepend`. Whereas the normal can_append (respectively can_prepend) returns True for *any* subtrajectory of a trajectory in the ensemble, the `strict` variant only returns True is the input trajectory is the beginning (respectively end) of a trajectory in the ensemble. This is useful when looking for a trajectory that satisfies the ensemble, such as when identifying a subtrajectories of a long trajectory that satisfy the ensemble. For the basic ensembles above, there is no distinction between these (in fact, there is also no distinction between can_append and can_prepend, since any trajectory that satisfies one must satisfy the other). However, for sequential ensembles, described in Sec. II E, there are significant differences, both for can_append vs can_prepend and for their `strict` and normal variants.

We stress that the basic formalism introduced here, connecting each path ensemble to a can-append criterion, is general and applicable beyond the ensembles implemented by OPS. For example, one could imagine an ensemble of all trajectories with an even number of frames, for which the corresponding can_append method would always returns True. OPS does not try to implement all possible ensembles; while the ensemble of all even-length trajectories could be implemented, it it not part of OPS due to its limited practical scientific use.

Note that the can-append criterion, as used by the shooting move (and similar trajectory generation approaches), results in what are called *candidate trajectories*. A candidate trajectory comes from the first trajectory that fails the can-append criterion. For some can-append criteria, such of that of the In_*X*_ ensemble, the can-append test ‘overshoots’, and only fails after the input trajectory could not possibly be in the desired ensemble. For others, such as that of the `LengthEnsemble`, can-append failure can be predicted before overshooting. To maximize efficiency, OPS trims the overshot frame to make candidate trajectories for ensembles where necessary, while not overshooting if not necessary.

### E. The sequential ensemble

#### 1. Definition of the sequential ensemble

One of the most productive ways to define useful ensembles in OPS is the `SequentialEnsemble`, which comprises a list of path ensembles that the trajectory must fulfill in the correct order. To understand this, consider a simple situation with a single state. Suppose we are interested in a path ensemble defined by a trajectory that begins in the state, then exits the state, then again returns to the state X. This ensemble can be summarized by the sequence [In_*X*_, Out_*X*_, In_*X*_]. Trajectories in this sequential ensemble can be split into subtrajectories that fulfill these three subensembles in the correct order.

Conceptually, a sequential ensemble consists of an ordered list of subensembles and an assignment algorithm to assign frames of a candidate trajectory to those subensembles. A trajectory satisfies the sequential ensemble if the assignment algorithm decomposes the trajectory into subtrajectories that satisfy each subensemble in the correct order. The can-append criterion forthe sequential ensemble can be defined based on the can-append criterion of the subensembles (and the assignment algorithm). While no unique choice for assignment algorithm exists, here we describe the approach used in OpenPathSampling.

A sequential ensemble is defined as an ordered set of (e.g., three) ensembles 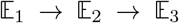 for which the in-dicatorfunction is

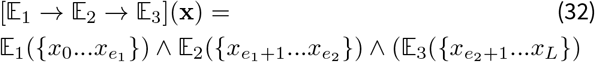

with frame indices *e*_1_ and *e*_2_ given by the assignment algorithm. Forthe assignment algorithm used in OPS, they are, respectively,

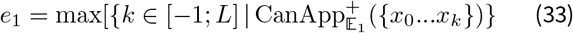

and

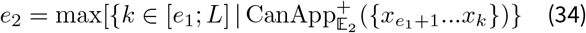

where the first equation Eq. 33 selects the maximum index *e*_1_ which still could fulfill the 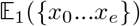 condition, and the second Eq. 34 likewise for *e*_2_. Note that here we make use of the fact that CanAppend returns True for an empty zero length trajectory to ensure that the index *e*_*i*_ always has a value. Naturally, the number of ensembles in the sequential ensemble can be arbitrary large:

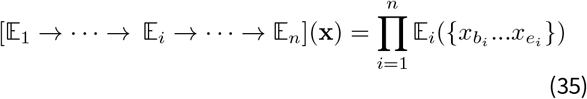

with *b*_*i*_ = *e*_*i*−1_ + 1, *e*_0_ = −1, and

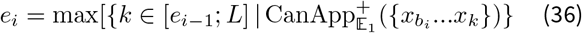

Note that if the first condition in Eq. 32 or Eq. 35 fails, all next conditions are not computed. The sequential ensemble thus is the set of trajectories that sequentially fulfill a set of ensembles.

The can-append criterion for the sequential ensemble is to use the frame assignment algorithm (the can-append of the subensembles) to assign all frames of the input trajectory to a subsequence of the subensembles. If all frames can be assigned to a subensemble, and if either (1) the subtra-iectory assigned to the last subensemble satisfies the can-append criterion for that subensemble, or (2) there are more subensembles later in the sequence, then the sequential ensemble’s can-append criterion is satisfied.

As an example of a sequential ensemble, considerthe situation with just two states *A* and *B* defined, and its union *S* = *A* ∪ *B*. The TPS ensemble connecting *A* and *B* can then be written as the sequential ensemble

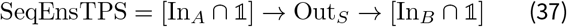

The indicatorfunction forthis ensemble SeqEnsTPS(x) returns True only if the first frame is in *A*, the last frame is in *B*, and no snapshot is in *A* nor *B* during the rest of the trajectory. Note that this function is identical to the *h*[x] in Eq. 5. A very similar expression is used forthe fixed length TPS:

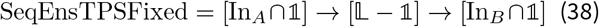

where the criterion is that the first and last slice are in A, and B, respectively, and the *L* − 1 slices are allowed to go anywhere.

The sequential ensemble forthe TIS ensemble is defined as

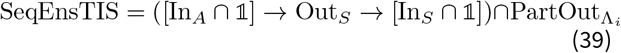

The corresponding indicator function SeqEnsTIS(x) returns 1 only if the first slice is in *A*, the last one ends in *A* or *B*, and no slice not in *A* nor *B* during the rest of the trajectory, but there is at least one slice that is not in the interface *i* volume. Note that this indicator function is identical to *h*̃_*i*_[x] in Eq. 7.

As a final example, the minus interface ensemble [25, 31] is

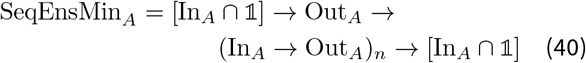

The indicator function for this ensemble SeqEnsMin_*A*_(x) returns unity if the first and last slice is in *A* and the trajectory leaves *A* at least once. Note that this definition allows multiple (*n*) entries into *A*. Here, we assumed that the boundary of A and the first interface are identical. Below, we discuss the OPS implementation of the minus ensemble for the more general case.

#### 2. Use of Sequential Ensembles for Path Sampling in OPS

The sequential ensemble is implemented in OPS by the `SequentialEnsemble` object. The test for whether a given trajectory satisfies the `SequentialEnsemble` uses the `strict_can_append` method of the underlying subensembles. It starts by making a candidate subtrajectory for the first subensemble, using that subensemble’s `strict_can_append` method until it returns `False`. The “`strict`” version is used because the subtrajectory that is assigned to the subensemble must satisfy the subensemble. If the resulting subtrajectory satisfies the first subensemble, then the process in continued with the next subensemble. This continues until no more frames can be assigned, either because all have been assigned or because there are no more subensembles. If all frames are assigned and all subensembles have been assigned a subtrajectory, then the given trajectory is in the `SequentialEnsemble`.

For most TPS/TIS purposes, one would like to stop integrating trajectories as soon as they enter the state. This is done by combining a volume ensemble, such as `AlllnXEnsemble(state)`, with a `LengthEnsemble(l)`, requiring an ensemble of length 1. This results in exactly one frame in the desired state. Hence, the `SeqEnsTPS` ensemble, Eq. 37, for which the initial and the final trajectory frame are in the initial and final states, respectively, but all other frames (at least one) are outside both states, is given by

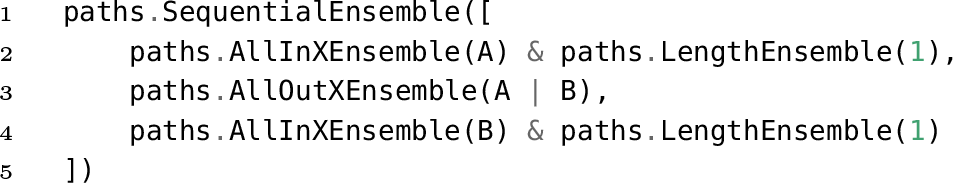

where we have made use of the set logic of the ensembles (see Eq. 29). The path should start with one frame in the initial state, then an arbitrary number outside either state, followed by precisely one frame inside the final state.

The slightly more complex sequential TIS ensemble SeqEnsTIS, Eq. 39, can be defined as

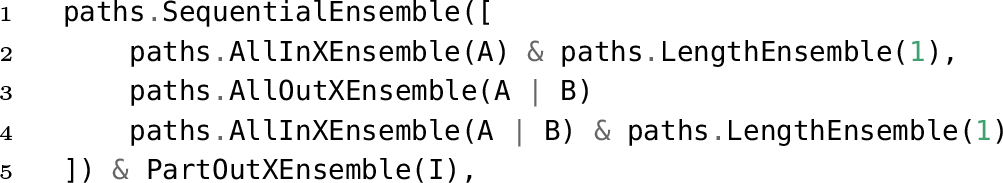

where *A*, *B*, and *I* are the volume-based definitions of state *A*, *B* and interface *I* respectively. Note that we have used set logic to define the union of *A* and *B* as a final state. Moreover, we require the middle part of the path to be outside of this union. Finally, the TIS ensemble requires at least a part of the entire trajectory to be outside the interface volume.

Fig. 2 provides an example, based on the TIS ensemble, of how frame assignment works fora sequential ensemble. Forsimplicity we leftoutthe interface crossing requirement. Two trajectories are shown, both of which fulfill the ensemble conditions. Each trajectory starts in *A*, which assigns the first frame in the (blue) 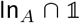 subensemble. Then there are a number of frames which are outside of the union of *A* and *B*, and which are assigned to the (black) Out_*A*∪*B*_ subensemble. The first frame that does not satisfy that criterion is also the last frame of each trajectory. Forthe top trajectory, the last frame is in *B*. Forthe bottom trajectory, the last from is in *A*. In both cases, the last frame satisfies the subensemble In_*A*∪*B*_.

**FIG. 2.**
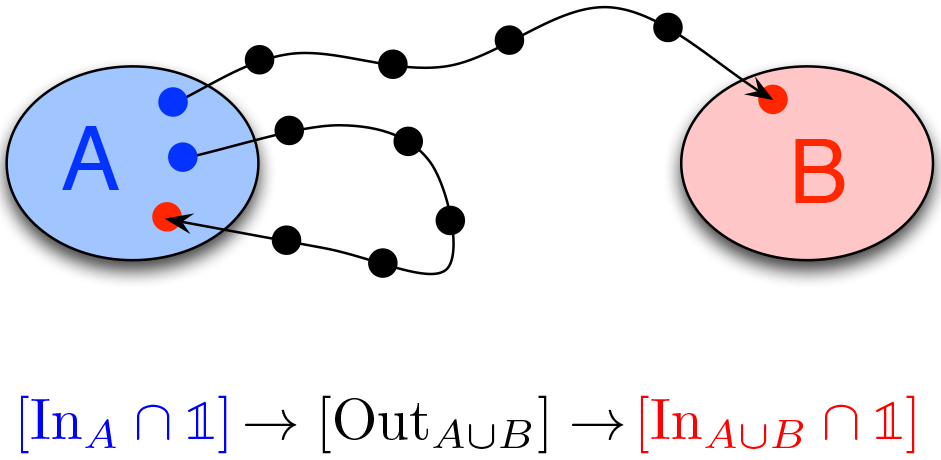
Frame assignment in an ensemble based on the TIS ensemble, for two trajectories. The points, which indicate individual frames in the trajectories, are colored to match the subensembles of the sequential ensemble, as given below the illustration. This ensemble differs from a real TIS ensemble because there is no interface.

In some cases, there is a need for an “optional” step in the sequence, which uses the so-called OptionalEnsemble. This means that a subtrajectory of a path can be in that ensemble, but does not have to be. In OPS this is implemented by forming the union of the ensemble 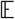 with a zero-length ensemble:

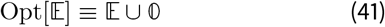

If no sub-trajectory fits the `OptionalEnsemble` a zero-length trajectory still allows the Sequential Ensemble to continue, thus effectively skipping the `OptionalEnsemble`.

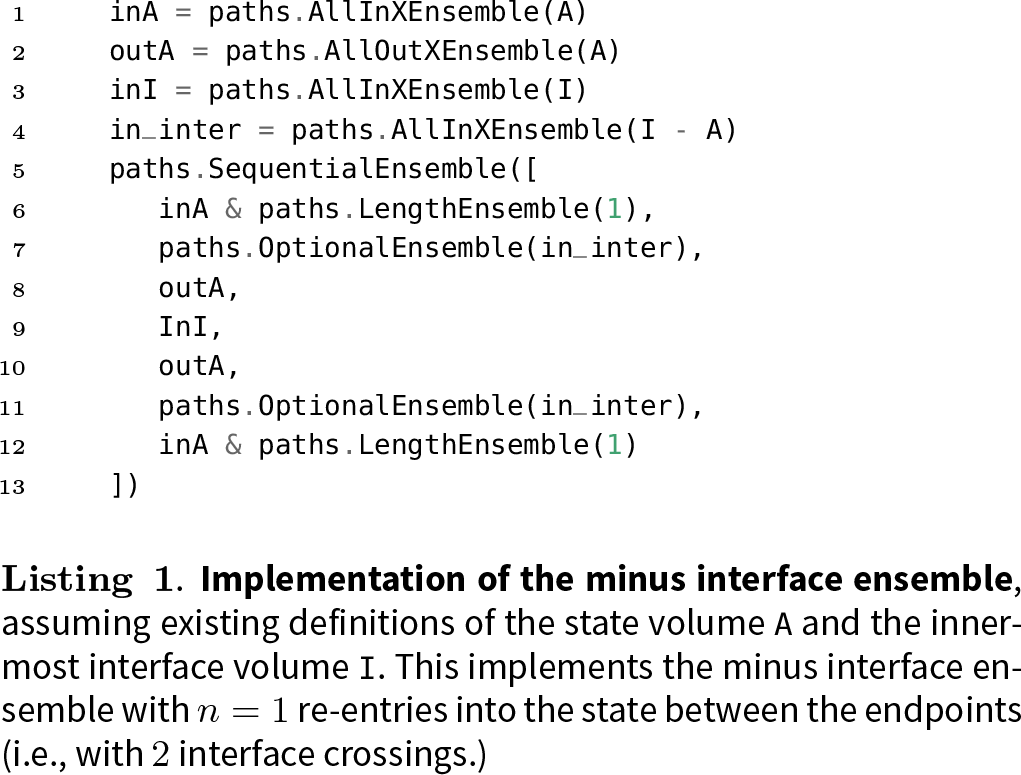

One example where we need to use the `OptionalEnsemble` class is when there is an interstitial space between the edge of the state and the innermost interface [30]. In simple cases, the innermost TIS interface *λ*_1_ is usually set to be exactly the boundary of the state *λ*_0_, but this is not required. A trajectory therefore can leave a state, visit the interstitial space and then cross the first interface, or it can skip this interstitial space in one frame and go directly from the state to the cross the interface. Both situations should be dealt with. The above TIS ensemble definition as already given works in this case. However, the minus interface ensemble needs special care. This ensemble is one of the more complicated ensembles in the TIS framework. As explained in Paper I [24], the minus interface ensemble is used (as part of the minus move) in RETIS to perform dynamics within the stable state, and return a new trajectory to one of the innermost TIS ensembles. This can be used to calculate the flux, to connect different interface sets in MISTIS, or to enhance decorrelation of trajectories [25,31]. The code forthe minus ensemble in Eq. 40 with *n* = 1 entries into *A* is given in Listing 1. Note the use of the `OptionalEnsemble` for the interstitial regions. This listing will also be discussed in more detail in following subsections.

#### 3. The reverse check and can-prepend for sequential ensembles

Up to this point, we have focused on sequential ensembles where the volumes associated with successive ensembles cannot overlap. That is, there can be no ambiguity as to which sub-ensemble a given frame of a trajectory is assigned to, regardless of the assignment algorithm used for the sequential ensemble. However, it is possible to define sequential ensembles where such overlaps are allowed, but these will become much more complicated and more subtle. In particular, special attention need to be paid to whether one can sample the same ensemble using the can_append and can_prepend methods.

OPS implements two main assignment rules. The normal OPS assignment algorithm is based on dynamics propagating forward in time (using 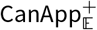) and, for clarity, can be called “forward assignment.” In the code, the forward assignment can be tested using `ensemble(trajectory)`. The alternative approach is based on dynamics propagating backward in time (using 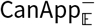), and will be called “reverse assignment”. The code to use it in OPS is `ensemble.check_ reverse(trajectory)`.

The reverse assignment algorithm is used to simplify the `can_prepend` 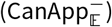 approach as implemented in OPS. The `can_prepend` algorithm for the OPS `SequentialEnsemble` is implemented analogously to the can_append algorithm. Both are “greedy” algorithms, in the sense that they try to assign the largest subtrajectory possible to the current subensemble. Since the forward assignment is greedy starting with the first subensemble of the sequential ensemble, and the reverse assignment is greedy starting with the last subensemble, the two algorithms might not yield equivalent results. Any ensemble that will be sampled with both forward and backward dynamics (as is done in the standard shooting moves in path sampling) must result in identical ensembles for both the forward assignment and the reverse assignment. Note that there are many cases in which the reverse assignment will not matter. For instance, generating initial trajectories (illustrated in Sec. IVA) or analyzing existing trajectories (see Sec. IV B) only require forward assignment. Moreover, many rare events methods (e.g. forward flux sampling [?]) involve propagating forward in time only.

Since the forward and reverse assignment algorithms are not equivalent, certain sequential ensembles could accept a trajectory when checked with the forward propagation, but not when checked with backward propagation (or vice-versa). For example, imagine volumes *A*, *I*_1_, *I*_2_ as illustrated in Fig. 3, where *I*_1_ ⊂ *I*_2_, and *A* ∩ *I*_2_ = ∅. Consider the ensemble 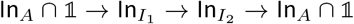. As shown in the top panel of Fig. 3, forthe illustrated trajectory, the forward assignment starts by assigning the first frame to the first ensemble ln_*A*_ (shaded blue in the figure). Then the frame assignment algorithm will look for a subtrajectory that satisfies the 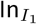 ensemble. In the example trajectory shown, it finds a two-frame subtrajectory that satisfies the 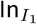 ensemble. The next frame is the first one that can be assigned to the 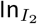 ensemble. Since frames that are in the volume *I*_1_ are also in the volume *I*_2_, the trajectory continues to assign frames to 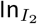 until it exits that volume and enters *A*. The last frame, in *A*, is assigned to the final 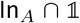 ensemble, shaded purple in the figure. Frames have been assigned to all ensembles, in the correct order, and no frames are unassigned. Therefore, this trajectory satisfies the ensemble.

**FIG. 3.**
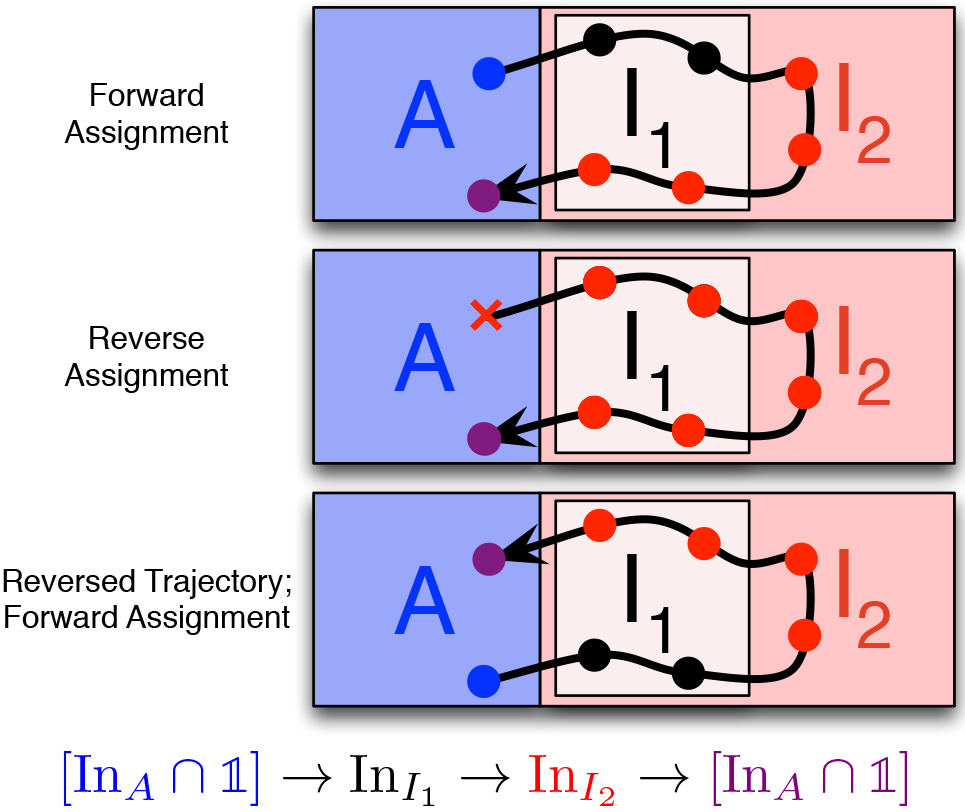
Applying different assignment approaches to a given trajectory. Points represent frames in the trajectory and are colored to show assignment, as with Fig. 2. The forward assignment algorithm (top) assigns frames in forward time order of the trajectory, with the forward order of the subensembles of the sequential ensemble. The reverse assignment algorithm (middle) assigns frames in the reverse order of the subensembles in the sequential ensemble, in reverse time order of the trajectory. Finally, the forward assignment of the reversed trajectory (bottom) is shown to illustrate that this is distinct from the reverse assignment. Note that a trajectory may be accepted by one assignment algorithm and rejected by the other (as shown here by the red X for an unassigned frame in the reverse assignment).

Next considerthe reverse assignment algorithm, as illustrated in the middle panel of Fig. 3. Assignment starts at the last frame of the trajectory, and at the final subensemble in the sequential ensemble. This frame is assigned to the final subensemble 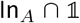 (shaded purple), as would also happen in case of forward assignment. Stepping backward along the trajectory, the assignment algorithm is looking for frames in the volume *I*_2_, following the penultimate subensemble. Since *I*_1_ ⊂ *I*_2_, it finds such frames, and it continues to find frames in *I*_2_ until the last frame to be assigned (the first frame of the trajectory), which is in *A*. Reaching that frame, the algorithm first checks whether it can be assigned to the 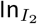 ensemble, as with the frame before. As this is not the case, the algorithm checks whether the frame can be assigned to the 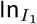 ensemble, the next subensemble in the reverse order. Since this is also not the case, the reverse check algorithm deems this trajectory to fail the sequential ensemble, as it does not contain subtrajectories assigned to all the correct ensembles in the correct order. In the figure, we signify this with the red x.

Note that the reverse assignment is not the same as using theforward assignmentalgorithm to assign framesfrom the time-reversed trajectory, as is shown in the bottom panel in Fig. 3. For this trajectory, the process of first reversing the trajectory and then assigning in the forward order leads to an assignment analogous to the forward assignment of the non-reversed trajectory. The trial trajectory will be accepted in this case.

The requirement for a trajectory to be sampled correctly with both forward and backward dynamics is that the forward and reverse assignment algorithms accept the same trajectories. For some sequential ensembles, such as the TPS and TIS ensembles, this means that the frame assignment is identical in both directions. However, this does not need to be the case, as can be seen from the minus ensemble implemented in Listing 1 and the trajectory assignments illustrated in Fig. 4.

**FIG. 4.**
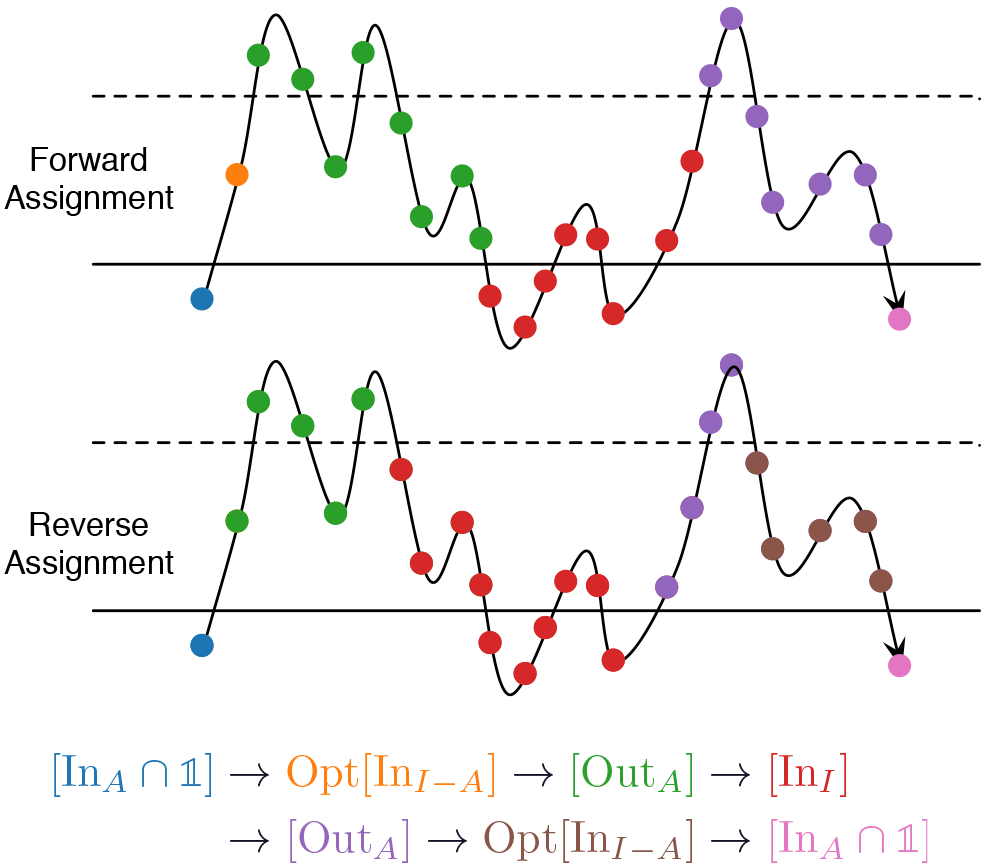
Frame assignment for an example trajectory in the minus ensemble. Points represent frames in the trajectory and are colored to show assignment, as with Fig. 2. The solid line represents the boundary of the state volume *A* and the dashed line represents the boundary of the interface volume *I*, with everything below the lines in the respective volume. The forward assignment algorithm (top) and reverse assignment algorithm (bottom) give the same result (the trajectory satisfies the ensemble), although the specific assignment of frames differs.

The minus ensemble includes trajectories that start with one frame in the state, go on to cross the interface, then return to the state, then cross the interface again, and finally end with one frame in the state. When the interface and state are not equivalent, there is an interstitial volume between them. This means that there could be recrossings of the interface or of the state boundary, as illustrated by the trajectory in Fig. 4, which also shows how this is handled by the careful implementation of the minus ensemble in Listing 1.

Recrossings are handled by using the fact that the criterion for failing a subensemble is to enter the next volume X. The not-yet-halt criterion for the subensemble is then the requirement to be in the complement volume 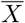, thus 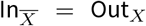. For instance, the green Out_*A*_ subensemble handles the condition that the trajectory should reen-terstate A, while allowing many crossings of the I interface. Likewise, the red 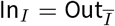 subensemble handles the second first exit of I, allowing many recrossings of the A boundary.

Note that some successive subensembles involve volumes that have overlap. For example, consider part of the minus ensemble Out_*A*_ → In_I_ → Out_*A*_. Because the associated volumes are not disjoint (i.e., *I* ∩ *Ā* ≠ ∅), frames in that intersection can be assigned to either ensemble, and will be assigned differently for the forward and reverse assignment algorithms. Additionally, the assignment of the optional ensembles depends on the assignment algorithm, again because of this volume overlap. The difference in the two assignment algorithms is shown for an example trajectory in Fig. 4.

Unlike the example in Fig. 3, any trajectory that satisfies the forward assignment for the minus ensemble will also satisfy the reverse assignment. Specific frames may be assigned to different subensembles, but the overall trajectory will either satisfy both assignment criteria or fail both.

In addition, because the minus move is one of the most computationally expensive moves in path sampling, we want to design the ensemble so that candidate trajectories are almost certain to be accepted. Without that requirement, the optional ensembles could be removed from the sequence and the Out_*A*_ ensembles in the sequence would become Out_*A*_ ∩ PartOut_*I*_. Here still, the forward and reverse frame assignments would differ. However, this would have the disadvantage that candidate trajectories could return to *A* immediately afterthe first frame in the interstitial, without crossing the interface. Such trajectories would be expensive to generateand would not beaccepted. Thesamplingwould be correct, but inefficient.

Because of the possible difference between frame assignment in theforward and reverse directions, it is important to know that the code defaults to forward propagation to check whether a trajectory is in an ensemble. We emphasize that an ensemble which does not give the same results in both directions can still be suitable for situations where only forward dynamics will be used (e.g.,generatingan initial trajectory), but will not suitable for approaches such as the shooting move in path sampling, which involves both forward and backward dynamics.

### F. Performance considerations

The previous sections provide a mathematically complete description of a new, set-based approach to describing path ensembles and their halting criteria in a consistent and unified manner. However, this approach, naively implemented, is not always computationally efficient. The functions described (such as the can-append criterion) take an entire trajectory as input, and therefore must iterate over all previously visited frames after each new frame is added. This leads to algorithms that scale as 𝒪(*L*^2^) instead of 𝒪(*L*) in *L*, the number of frames. In OPS, this scaling problem is managed by using caches for the sequential ensemble, combined with a Boolean parameter trusted that can be passed to the can_append and `can_prepend` functions (as well as their `strict` variants). The `trusted` parameter for can_append indicates that, as of the previous frame, the trajectory satisfied the can_append criterion (and similarly for `can_prepend` and the `strict` variants). Additionally, the ensemble indicator function, given by `ensemble(trajectory)`, takes an optional Boolean parameter called candidate. If `candidate=True`, then the code assumes that the trajectory was generated by the `can_append` or `can_prepend` method, and therefore only certain parts of the trajectory need to be tested.

For example, consider a flexible-length TPS ensemble as in Eq. 37 and a trajectory (*x*_0_,…, *x*_*i*_). If (*x*_0_,…, *x*_*i*−1_) satisfied the can-append criteria, then we know that the last trajectory with frames assigned was Out_*S*_, and we should first check whether *x*_*i*_ ∉ *S*, which would allow us to assign it to that subensemble as well. The trusted parameter tells us that we can trust that the previous frame passed can-append, enabling a faster path for checking the can-append criterion. In addition, the `SequentialEnsemble` keeps a cache of the frame assignment, so the algorithm knows immediately to which ensemble the frame should be assigned (with safety checks that this frame is still part of the same trajectory.)

As an example of the use of the candidate parameter, again consider the flexible-length TPS ensemble, with some trajectory (*x*_0_,…,*x*_*L*_). If that trajectory was generated using the `can_append` or `can_prepend` rules, no frames except the first and last can be in any state. In this case, we can check whether the trajectory satisfies the ensemble just by checking if the first and last frames are in the appropriate states. The methods built into OPS for arbitrary ensembles are general, but might not be the most efficient. Customized ensembles can make use of the trusted and candidate parameters to provide faster calculations for trajectories known to be generated by dynamics, while still benefitting from the general approaches for trajectories of unknown origin.

### G. Guidelines for designing custom ensembles

The above sections introduced the set-based notation for path ensembles, illustrated the connection between this notation and the innerworkings of OPS, and showed howOPS uses this conceptual framework to implement ensembles used in path sampling simulations. In Sec. IV, we will provide several more examples of useful path ensembles. However, defining new ensembles might not seem completely straightforward. To help bridge the gap between understanding the ensembles we present, and creating new ensembles, in this section we provide some general guidelines and tricks that could be seen as rules of thumb for ensemble building.

#### • Use anchors

In many path ensembles, trajectories start and end with a frame in a specific volume (or union of volumes). It can be useful to think of these as anchors to start designing the ensemble. Typically, the building block is a single frame in some volume *A*, i.e., the ensemble 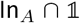.

#### • Use complement ensembles and volumes

If we want a trajectory to end with the first frame in some volume *A*, we might think of this as a PartIn_*A*_ ensemble. But, as discussed in Sec. II D, the PartIn_*A*_ ensemble never halts. However, the first trajectory that will satisfy it can come from the first trajectory that does not satisfy its complementary ensemble, Out_*A*_. This can, of course, also be written as In_*Ā*_. In some cases, the complement volume may be the one that is more naturally defined. For example, if part of a sequential ensemble is supposed to lead to a first frame in *A*, we can use the Out_*A*_ ensemble, as is done in the minus ensemble, which also uses this to create a first frame outside of the interface *I* using In_*I*_ (where it is more natural to refer to a frame outside the interface volume than a frame inside the complement of the interface volume). A more elaborate example using this reasoning is given in Sec. IV for the ensemble used to get the initial trajectory that visits all states.

#### • Think in terms of halting criteria

First, it is important to make sure that every ensemble (and every subensemble of a sequential ensemble) will eventually halt. Second, the halting criteria can be useful when designing sequential ensembles. Because the sequential ensemble uses a greedy algorithm, it is important to think in terms of the stopping criteria of the previous ensemble and where that leaves you. The previous guideline explained how to get a crossing out of some volume (call it *I*), but typically we speak of crossing from starting in one volume (call it *A*) and then exiting the volume *I*. To know which volume the subensemble will start in, look at the previous subensemble and apply the rules in the guideline about complement ensembles and volumes.

#### • Prefer set logic on volumes, not ensembles

When creating ensembles, set logic on ensembles and on volumes might seem very similar. For instance, one might be tempted to code the ensemble Out_*A*∪*B*_ as Out_*A*_ ∪ Out_*B*_ However, this is incorrect, as explained in Section II C. Also, the logical complement of ensembles in general is not what one naively would expect. The set logic for volumes is usually more familiar, and therefore, whenever possible, should be used.

#### • Beware of unions with PartIn or PartOut

This relates to both the suggestion of thinking in terms of halting criteria and preferring set logic on volumes. The danger here is that, while an ensemble such as In_*I*_ ∪ PartOut_*A*_ may seem reasonable, the stopping criterion of PartOut_*A*_ is to never stop, and therefore a union with it leads to infinite trajectories.

#### • Check the reverse assignment

If using path sampling, or any other algorithm that requires generating dynamics in the backward time direction, check that the reverse assignment gives the same results as the forward assignment. The ideas behind this are described in Sec. II E3. Developing a symmetric sequence for the sequential ensemble can help with the reverse assignment. Not all methods require that the reverse assignment be used; only those that involve propagating backward in time.

#### • Use optional ensembles for flexibility

The optional ensemble allows a particular subensemble of a sequential ensemble to be skipped. This is useful when the ensemble will be employed in many variants, and when it is uncertain whether the subensemble will always be satisfied (as with the minus ensemble). Including optional ensemble can also ensure that both the forward and reverse assignment work properly.

#### • Use unions of complex ensembles

Sometime a union of complex ensembles, such as sequential ensembles, is the best way to achieve a desired ensemble. For example, consider sampling *A* → *B* transitions and *B* → *A* transitions in one ensemble. A TPS ensemble from *A* ∪ *B* to *A* ∪ *B* will not work, since it will allow *A* → *A* and *B* → *B* transitions. Instead use a union of sequential ensembles, SeqEnsTPS_*A*→*B*_ ∪ SeqEnsTPS_*B*→*A*_.

Finally, there is often more than one way to implement a given path ensemble. Theseguidelinesshould providetools both for the design of new ensembles, as well as to understand ensembles we have provided as examples in Sec. IV.

## III. GENERAL FRAMEWORK FOR CUSTOM MONTE CARLO APPROACHES

Transition path sampling amounts to Monte Carlo of trajectory space. In standard TPS, the Monte Carlo procedure samples trajectories from a single path ensemble. In TIS, and particularly in RETIS, the Monte Carlo procedure simultaneously samples trajectories from an expanded ensemble, combining multiple interface ensembles. Standard TPS can be seen as a special case of this expanded ensemble, where only a single path ensemble is sampled. The expanded ensemble gives rise to the `SampleSet` object in OPS, as described in Paper I.

Monte Carlo moves in OPS, based on this expanded ensemble, are performed by the PathMover object. PathMovers can change trajectories within the ensemble being sampled (as with the shooting move), or they can alter the ensembles associated with trajectories without changing the trajectories (as with path replica exchange), or they can alter both the trajectory and its associated ensemble (as with the minus move).

The PathMovers are organized in an overall *move decision tree.* This tree includes, besides the movers that perform the Monte Carlo steps, several so-called *structural movers*, e.g., the `RandomChoiceMover` that randomly selects one of several *submovers* (these structural movers are described in Sec. III A). In principle, a manually assembled move decision tree is all that is necessary for a path sampling simulation. However, for complicated move decision trees, this becomes tedious and difficult. The PathMovers, including the structural movers, constitute a low-level interface that is sufficient, but not particularly user-friendly. Therefore, we have developed a higher-level layer, usingthe `MoveStrategy` and `MoveScheme` objects, which automates the repetitive lower-level operations, and enables the user to customize the move decision tree easily.

The `MoveScheme` is an overall container that builds the move decision tree, while each `MoveStrategy` deals with a part of that tree: providing, for example, details on how the shooting move will be performed, or which pairs of ensembles are involved in replica exchange. A path sampling simulation will have one `MoveScheme`, and that `MoveScheme` will include multiple `MoveStrategy` objects.

The following subsections describe how the structural path movers allow combining existing movers into a more complicated move, and how to use the `MoveStrategy` and `MoveScheme`. Subclassing existing objects can create more complicated path movers; details are available in the online documentation for OpenPathSampling at http://openpathsampling.org.

### A. Structural Movers

`PathMover` objects such as the `OneWayShootingMover` and `ReplicaExchangeMover` generate newtrial paths. However, they need to be organized into the overall move decision tree. This organization is done by other subclasses of PathMover, which we call *structural movers.* Important structural movers include:

- A `RandomChoiceMover`, one of the main structural elements in most move decision trees, randomly selects one of its submovers. For example, a first `RandomChoiceMover` selects the type of move (shooting, replica exchange, etc.), followed by a second `RandomChoiceMover` that selects a mover for the specific ensemble(s) involved in the move. The `RandomChoiceMover` is also an important element in many PathMovers. For example, the `OneWayShootingMover` consists of a `RandomChoiceMover` that selects between a `ForwardShootingMover` and a `BackwardShootingMover`. By default, the submovers within a `RandomChoiceMover` have identical probability of being selected; this can be changed with the weights parameter at initialization.
- A `SequentialMover` employs several submovers in a specific order, where each submover is accepted independently. This mover is not a combined trial move, but a bundle of several moves together in a specific order.
- The `ConditionalSequentialMover` is similar to a `SequentialMover`, but provides an early-rejection scheme, which is important for moves where a failure in an early step can guarantee that the whole move fails, especially if later steps are very expensive. Below we will discuss how this plays a role in the `MinusMover`.
- An `EnsembleFilterMover` removes resulting Samples associated with intermediate ensembles from the results. In complicated movers, extra, internally-defined Ensemble objects can create intermediate steps in the mover, which would end up in the results. The `EnsembleFilterMover` filters those (often uninformative data) out of the results.

The move decision tree can take many forms. To obtain information about the path mover most likely of interest (e.g., `ReplicaExchangeMover` or `ForwardShootingMover`) regardless of the specific structure of the move decision tree, we implemented a property canonical in the MoveChange. As discussed in Paper I[24], a `PathMover` takes a `SampleSet` as input, and returns a MoveChange object. This MoveChange can contain other MoveChanges from submovers; in this way the whole path taken through the move decision tree is preserved. However, the nested structure of MoveChanges can make it cumbersome to access attributes from the MoveChange of the specific submover of interest (e.g., shooting point from the MoveChange associated with the shooting mover). Therefore, the MoveChange.canonical property directly accesses the first nested MoveChange associated with a mover that identifies itself as “canonical” Subclasses of `PathMover` can declare themselves canonical by setting the class attribute `_is_canonical` to True. Examples using the canonical property can be found in Paper I, Sec. VI.A 6.

The `MinusMover` provides a useful example of how several structural movers can be put together to generate a new Monte Carlo move. As described in paper I, the OPS `MinusMover` is, in a way, a combination of replica exchange and shooting moves. In MSTIS, each state typically has one MinusMover, which takes trajectories from two ensembles as input: the TIS innermost interface ensemble and the minus ensemble. In the discussion that follows, the state is denoted A and the innermost interface volume *X*. In many cases the state definition A will be identical to *X*, but this is not required.

Both the TIS ensemble and the minus interface ensemble are described in Sec. II E. In the minus move, both use the same interface volume, *X*. In addition, there is an ensemble which is used internally in the minus move. This ensemble is nearly the same as the TIS ensemble used as input, except that instead of allowing paths to end in either *A* or make a transition to another state *B*, all paths in this internal ensemble must start in *A*, cross the interface, and also end in *A*, i.e., this is the ensemble 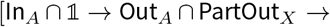 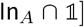. In the context of the minus move, we refer to this as the “segment ensemble" Trajectories in the minus interface ensemble begin and end with subtrajectories that also satisfy the segment ensemble and therefore satisfy the innermost interface ensemble.

The minus move consists of three steps: (1) randomly selecting one of the subtrajectories that satisfies the segment ensemble from the minus ensemble trajectory; (2) performing a replica exchange move between the selected segment and the path in the innermost interface; (3) extending that path in a random time direction until it satisfies the minus interface ensemble.

The first step is performed by a `RandomChoiceMover` that selects between eithera `FirstSubtrajectorySelectMover` or a `FinalSubtrajectorySelectMover`, where in both cases the subtrajectory satisfies the segment ensemble This step should always be accepted as the initial path satisfies the minus ensemble.

When there is only one innermost ensemble, the second step is just a replica exchange. This replica exchange can only fail if the innermost interface path happens to cross to another stable state. In the case of a multiple interface sets (as in MISTIS), however, there are multiple innermost interfaces [31]. The interface to exchange with is selected with a `RandomChoiceMover`that includes `ReplicaExchangeMovers` for each innermost interface. The segment might not overlap with the selected interface, and therefore this step very well mightfailforMISTIS.

In the final step, the trajectory that was initially in the innermost interface ensemble is extended until it satisfies the minus ensemble, using a `RandomChoiceMover` to choose either a `ForwardExtendMover` or a `BackwardExtendMover`.

No part of the move can be accepted unless all parts succeed. Therefore, we use a `ConditionalSequentialMover` for this. In addition, we used the intermediate “segment” ensemble. To remove this from the results, we wrap the mover in a `EnsembleFilterMover`.

Fig. 5 shows the internal structure of this mover. Normally, this structure is not shown in the move decision tree visualization because the `MinusMover` is marked as a canonical mover, and the visualizer does not show internal structure of canonical movers. This setting can be overridden by changing the `options.analysis` dictionary of the visualizer.

**FIG. 5.**
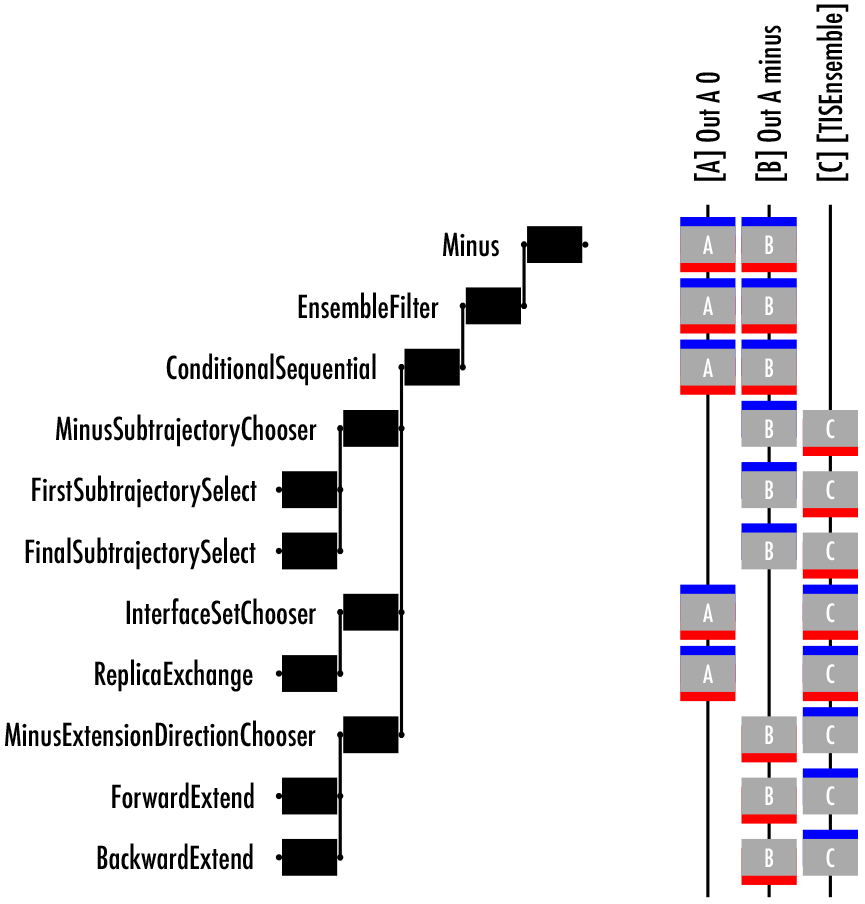
Internal structure of a MinusMover. At the left, the hierarchical structure of mover. Each layer (moving left) is encapsulated in the objects to the right. During a minus move, submovers are visited from top to bottom (but not all submovers are visited). The `MinusMover`, labelled “`Minus`”, is the outermost container. Inside it is the `EnsembleFilterMover` that filters out results in the internally-used segment ensemble. The primary sequence of the minus move is in the `ConditionalSequentialMover`, each of which involves a `RandomChoiceMover`, which select the specific movers to do at each stage. On the right are the three ensembles involved in the minus move, labelled with A, B, and C. Ensemble A is the innermost TIS ensemble, ensemble B is the minus ensemble, and ensemble C is the segment ensemble. The input and output ensembles for each mover are shown in blue and red (respectively). Movers like replica exchange have the same input and output ensembles (even if the replicas have changed), whereas movers like the subtrajectory selectors move a replica from one ensemble to another.

### B. MoveScheme and MoveStrategy objects

The `MoveScheme` contains multiple `MoveStrategy` objects, used to automatically build the (often elaborate) move decision tree. It also is possible to build move decision trees manually. To make these manually-built trees compatible with other parts of OPS, the root mover needs to be wrapped in a in a `LockedMoveScheme`. However, many of the tools for `MoveSchemes` are not available for `LockedMoveSchemes`, and much of what is discussed in the remainder ofthis section is not applicable to `LockedMoveSchemes`.

In addition to containing the move decision tree, a `MoveScheme` organizes information about the sampling process, and provides access to tools for analyzing the sampling procedure after the simulation. In particular, the `MoveScheme` organizes the movers into labeled or named groups. For the default TIS scheme, the group names are ‘`shooting`’ (shooting movers), ‘`repex`’ (replica exchange), ‘`pathreversal`’ (path reversal), and ‘`minus`’ (minus move). These can be accessed with `scheme.movers[group_name]`. Each group consists of multiple `PathMovers`. Each `PathMover` in turn has specific input and output ensembles. For shooting, a single ensemble is used for both input and output. For `EnsembleHopMovers` (the single-replica version of replica exchange, which moves a single replica from one ensemble to a different ensemble), the input mover is different from the output mover. `ReplicaExchangeMovers` have two input and two output ensembles (the same pair for both). We refer to this set of input and output ensembles as the mover’s *ensemble signature*.

To build the move decision tree, one must decide (1) which ensemble signatures will be part of each type of move (e.g., which ensembles to shoot from), (2) how the movers will be implemented (e.g., what kind of shooting move to use), and (3) how all movers are organized into the overall decision tree (e.g., select type of mover first, then specific ensembles to include). `MoveScheme` builds the move decision tree by applying multiple `MoveStrategy` objects. Each `MoveStrategy` is associated with a specific priority level. When building the move decision tree, the strategies are applied in order of priority level, and within a priority level, in order of addition to the `MoveScheme`. The default priority levels are in `openpathsampling.strategies.levels.LEVELNAME`, where `LEVELNAME` can be, in order of ascending priority, `SIGNATURE, MOVER, GROUP, or GLOBAL`. Internally, these priority levels are represented by integers between 0 and 100, but these specific levels are named for convenience. The recommended use for these levels is as follows:

- Strategies at the `SIGNATURE` level modify the set of ensemble signatures to be used; for example, the default `NearestNeighborRepExStrategy` which executes replica swaps among neighboring interfaces, versus the `AllSetRepExStrategy`, which implements replica swaps among all interfaces within the same `Transition` (interface set).
- Strategies at the `MOVER` level customize the behavior of the created path movers; for example, the `OneWayShootingStrategy` provides the ability to choose a shooting point selector for the shooting group.
- Strategies at the `GROUP` level can take the already-built movers and rearrange them, or convert them to a different approach. For example, the single-replica EnsembleHop movers can be converted to normal replica exchange using the `ReplicaExchangeStrategy`, which is a GROUP-level strategy. In Sec. IV, we show another GROUP-level strategy, which will take the already-created shooting and replica exchange movers, and reorganize them into a single sequential move.
- Strategies at the `GLOBAL` level organize the overall move decision tree. The two `GLOBAL`-level strategies in OPS are the `OrganizeByMoveGroupStrategy`, which first selects a move type, and then selects which specific mover (which signature, i.e., which ensembles), and the `OrganizeByEnsembleStrategy`, which starts with the selection of the ensemble to move (as necessary with single-replica approaches) and then selects which type of move to do.

By using this priority-level system, the move decision tree can be built correctly, regardless of the order in which the specific move strategies are added to the scheme. Parts that must be built later *are* built later because of the priority levels. In addition, the details of what has already been built (e.g., specific choices of parameters for movers, such as the shooting point selection in a shooting move) can be retained by later moves that reorganize the entire move decision tree.

`MoveStrategy` objects deal with the details of the implementation, when building the move decision tree. For example, two-way shooting could be used instead of one-way shooting by starting with the default scheme applying a `TwoWayShootingStrategy` to the scheme:

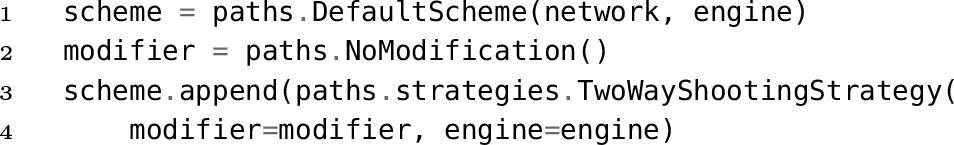

where `engine` and `network` were already appropriately defined (see Paper I for details[24]). This implements a two-way shooting move decision tree where the shooting point is not modified, and the decorrelation depends on the stochastic dynamics. Alternatively, for NVT dynamics, one could choose the momenta randomly from a Maxwell-Boltzmann distribution, as is, for instance, done in the aimless shooting [32] approach. The scheme then becomes

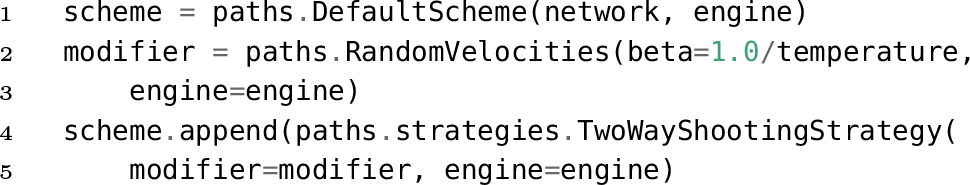

where `temperature`, `engine`, and `network` were already appropriately defined. This implements a two-sway shooting move decision tree where the shooting point obtains random velocities taken from a Maxwell-Boltzmann distribution at the desired temperature. If the engine supports constraints, the `RandomVelocities` modifier will project that distribution into the space of constraints.

As another example, a modified shooting point selector could be used, instead of the default (uniform selector). For instance, the `GaussianBiasSelector` selects the shooting point according to a Gaussian probability in a specified collective variable [33]. After setting up the `scheme` as above, and creating a collective variable cv, the following code will implement this selector:

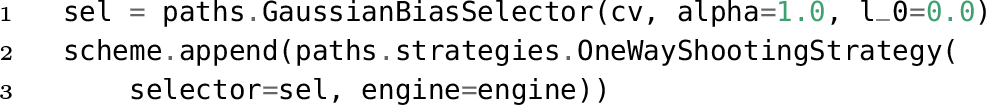

where the Gaussian bias is defined by exp(−*α*(cv(*x*) − *l*_0_)^2^), with *x* a snapshot, and *l*_0_ and *α*, respectively, control the position and width of the Gaussian.

Athird example of a `MoveStrategy` is to add a specific ensemble pair to the list of possible replica exchange moves. To do this, one would first select the ensembles (call them `ens1` and `ens2`). Then, after creating the `scheme` as before, the new replica exchange pairs can be added with:

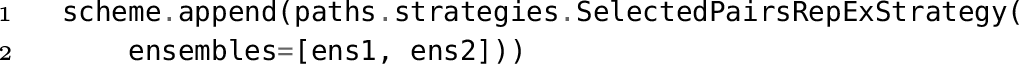

Note that this last example was a signature-level strategy, whereas the other examples were mover-level. The priority-level system used in OPS means that the user does not have to consider the order in which the strategies are built when appending them to a custom move scheme. For example, the signature-level strategy that changes the ensembles used in replica exchange can be appended afterthe mover-level strategy that provides details of how to perform the shooting move, even though the order they are used in the opposite orderwhen buildingthe move decision tree.

## IV. ILLUSTRATIVE EXAMPLES

### A. Creating new ensembles

One of the significant features of `OpenPathSampling` is the ability to generate valid paths for arbitrary path ensembles. This capability facilitates the development of new methodologies, which often require the creation of new path ensembles. In addition, this feature has practical advantages for users as well. In the Appendix of Paper I, we briefly mentioned one such practical use[24]. To obtain the initial conditionsfora path samplingsimulation, we can use a high temperature trajectory. For multiple state TIS simulations, we also need initial trajectories that satisfy all the ensembles in the MSTIS network: there must be trajectories that begin in each state, and which exit each interface volume. For most MSTIS simulation setups, a path that undergoes a transition to another state, crosses all the interfaces associated with the starting state. Therefore, we can use such a transition path as an initial path for all interfaces. And since we can reverse paths, a long trajectory that visits all states will contain, for each of the defined MSTIS ensemble in the network, a (possibly reversed) subtrajectory that satisfies that ensemble.

Thus, our high temperature target trajectory is one that has visited all states. To define an ensemble that will generate such a trajectory, we use complement ensembles and think in terms of halting criteria, as suggested in the guidelines in Sec. II G. We need a condition that remains true until the trajectory has visited all states — in other words, the opposite of the condition that the trajectory has visited all states. This means that we should continue as long as the trajectory has not visited at least one state. We can express this idea as a path ensemble

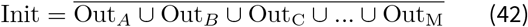

which is only true if none of the *M* Out conditions are fulfilled. The continuation condition is now the negation of this ensemble

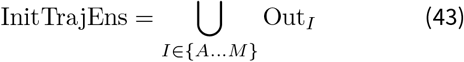

This condition translates in OPS to the python code

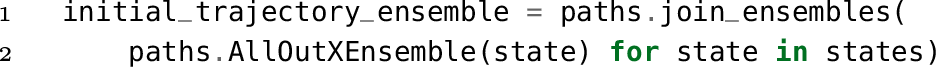

where `states` is a list of the state volumes. We can then create the goal trajectory using the `engine.generate` method by

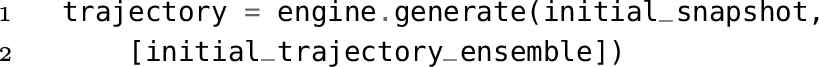

The resulting `trajectory` will have visited every state, and the last frame will be in the last state visited. Since it visits every state, then it has, for every state, a subtrajectory that starts in that state and ends in another (in some cases requiring time reversal). This means that subtrajectories of this long trajectory can be found to satisfy all the ensembles in the MSTIS network. Note that we use the ensemble itself asa condition, notitsCanAppfunction. Inthiscase, because these are In/Out_*A*_-type ensembles, the CanApp is equivalent to the ensemble check (this is not necessarily the case for other ensembles). Conceptually, we are after the first trajectory that does not satisfy the ensemble, so we use the ensemble check itself.

We can also define another (arbitrary) ensemble to obtain a first trajectory suitable for the TIS bootstrapping procedure. This procedure takes a trajectory satisfying the innermost interface ensemble of a TIS transition, and performs shooting moves until the resulting paths satisfy the ensemble(s) forthe subsequent interface(s). To get that initial trajectory, we want to start from any arbitrary frame, then have at least one frame in state *A*, then have at least one frame that crosses the interface, and end with exactly one frame in either state *A* or state *B* (where state *B* can be generalized to the union of multiple other states). The sequential ensemble to do this is

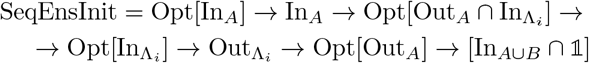

Following Sec. II G, this ensemble uses an anchorthat combines the optional ensemble outside of *A* with a required ensemble inside *A*. As also suggested in the guidelines, the `OptionalEnsembles` in this ensemble are designed to ensure that any possible trajectory that satisfies the overall goal will still be accepted. This ensemble translates into OPS code as

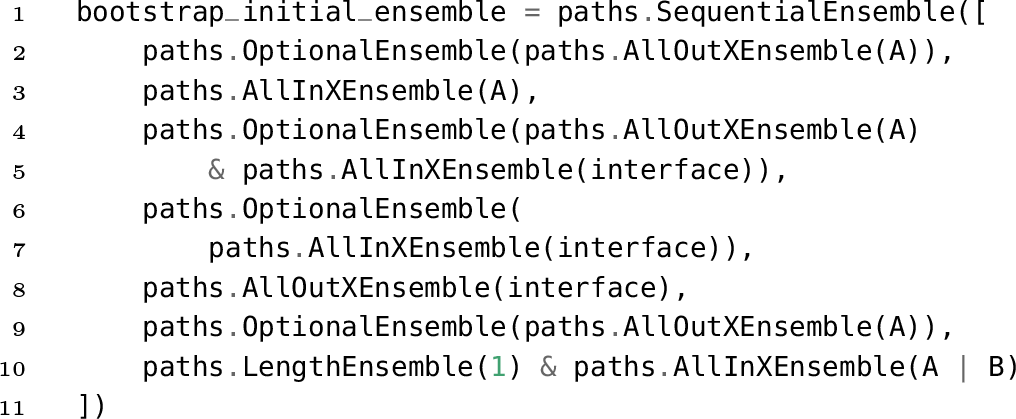

A similar ensemble is part of the `FullBootstrapping` calculation, which fills TIS ensembles starting from a single snapshot.

### B. Using Ensemble.split for trajectory analysis

The `Ensemble` object in `OpenPathSampling` provides a convenient way of analyzing trajectories in terms of subtrajectories. The `ensemble.split(trajectory, overlap)` method takesa long `trajectory` and returnsa list of subtrajectories that satisfy the `ensemble`. Successive subtrajectories will have at most `overlap` frames in common (with a default of 1 shared frame). For example, we can check whether any trajectories in a fixed-length TPS simulation included recrossings. Given state volumes A and B, we first create the *B* → *A* transition ensemble:

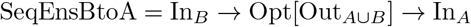

which in OPS code translates to

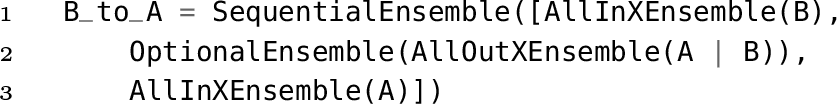

Making frames outside of both A and B optional captures trajectories where the transition occurs without any intermediate frames (this is unlikely in our examples, but could be common in long paths with (too) infrequent saving of frames). We can apply `B_to_A.split(trajectory)` to every trajectory accepted bythefixed-lengthTPSensemble. Ifthe resulting list is not empty, a recrossing in the *B* → *A* direction took place. Since the first frame of every accepted trajectory has to be in *A* and the last frame of every trajectory is necessarily in *B*, the existence of a *B* → *A* transition guarantees a recrossing. Forthe fixed path length alanine dipeptide example from Paper I[24], Sec. VI A, we find 109 accepted trials with recrossings, including 5 with 2 recrossing events. For accepted paths with a single recrossing, there are two *α* → *β* transitions in the path – one before recrossing, and one after. With two recrossings, there would be three transitions. An example of an accepted recrossing trajectory is shown in Fig. 6.

**FIG. 6.**
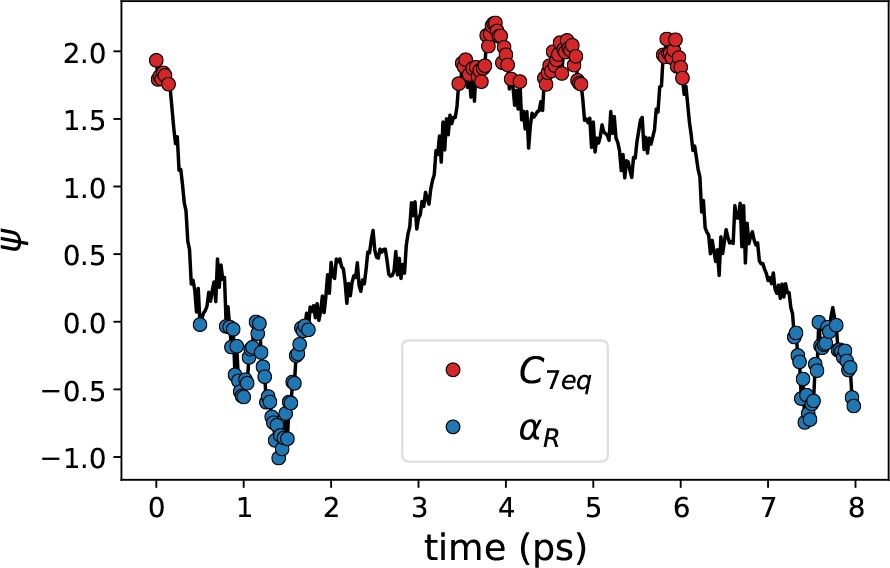
**Accepted TPS trajectory with recrossing** taken from the fixed-length TPS simulation of alanine dipeptide in Paper 1[24]. The angle *ψ* is plotted as a function of time. Frames in 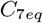 are marked in red; frames in *α*_*R*_ are marked in blue.

This approach also allows distinguishing between multiple channels fora given reaction. For instance, we can compare the behavior of fixed path length TPS and flexible path length TPS for alanine dipeptide. By taking `ensemble` as the flexible path length TPS ensemble, application of the split function identifies subtrajectories of the fixed path length TPS that match the flexible path length ensemble.

Fig. 7a shows path-length histograms forthe transitions in the flexible path length TPS ensemble and in the fixed path length ensemble (selected using the `split` function), from our TPS simulations of alanine dipeptide reported in Paper I. These histograms differ because the fixed length ensemble in fact sampled two different transition mechanisms. To show this, we can define custom path ensembles that distinguish between the two mechanisms. First, we define additional volumes, based on the *ψ* collective variable: an A volume for 100 < *ψ* < 200 (where the `CVPeriodicRangeVolume` automatically wraps into the correct bounds), and a B volume for −100 < *ψ* < 200. These two volumes are based on the original states, but have no restrictions in *ϕ*. Two additional volumes account forthe “noman’s land” region outside the states: `nml_increasing` for −160 < *ψ* < −100, and `nml_decreasing` for 0 < *ψ* < 100. Next, we identify a transition as“increasing” ifthe trajectory crosses the `nml_increasing` volume (i.e., the value of 0 increases while going from one state to the next), and “decreasing” if it crosses the `nml_decreasing` volume. The sequential ensemble forthe increasing transitions can be defined by

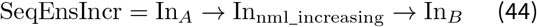

which is in OPS code

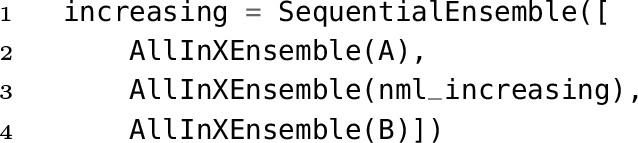

**FIG. 7.**
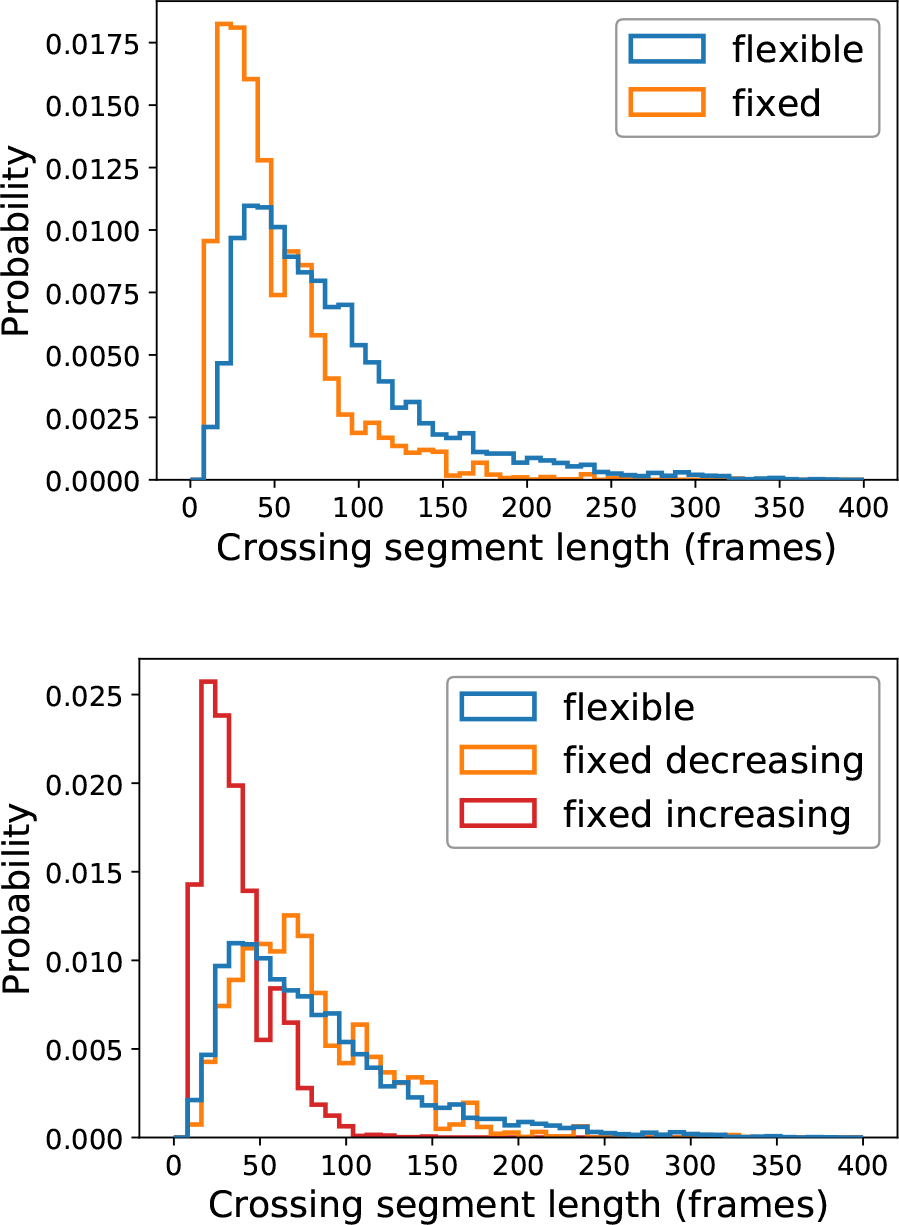
Transition path length distributions. Top: Without distinguishing between increasing and decreasing transition types from the fixed length TPS. Bottom: Distinguishing the increasing and decreasing transitions. The difference in the path length distributions can be attributed to the fixed-length simulation sampling both increasing and decreasing transitions.

The ensemble for decreasing transitions is defined similarly, but using nml_decreasing in place of `nml_increasing`. Trajectories that satisfy the original TPS ensemble will have subtrajectories that satisfy one of these ensembles. By using the `ensemble.split()` method, we can identify which mechanism each trajectory represents.

Fig 8 shows several trajectories (blue and purple) connecting the original two stable states (dark red). The states are defined in terms of periodic variables, and can wrap around the periodic boundary. The “extended” versions of the states that were defined above are shown in light red, and the two different “no man’s land” volumes are labelled nml_decreasing and nmljncreasing. Several hypothetical trajectories are shown, all of which would count as a decreasing transition. Trajectory (a) would be easy to analyze by a volume-based approach (and is also a realistic trajectory for this system). Trajectories (b) and (c) would count as decreasing transitions, but an analysis based only on volumes (without consideration of ordering) might miss them. Trajectory (d) includes both an increasing and a decreasing transition (and would be extraordinarily unlikely in this system).

**FIG. 8.**
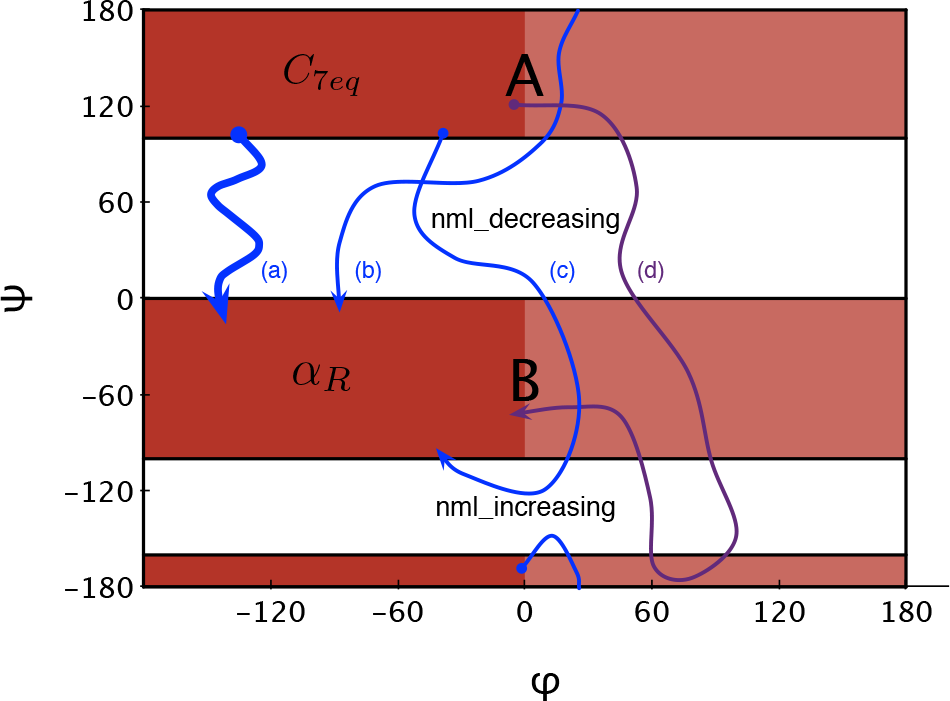
Volumes and example trajectories for the different ways a “decreasing” transition can occur. The shaded areas represent the extended state definitions used in the analysis of the transitions, while the darker shaded areas are the actual states of the system, (a) A typical decreasing path, (b) and (c) Paths which visit the “increasing” no man’s land, but only transition across the decreasing, (d) Path with both increasing and decreasing transitions.

The fixed length alanine dipeptide example shows 3570 trajectories in the decreasing channel and 6670 trajectories in the increasing channel (weighted by the Monte Carlo weights from the TPS simulation). All transition trajectories satisfy exactly one of the two channels, although some TPS trajectories, due to recrossings, have more than one transition trajectory. The flexible length example has all its trajectories in the decreasing channel. The existence of recrossings in the fixed length TPS ensemble demonstrates that these transitions in alanine dipeptide are not *that* rare, so it is not surprising that we would also observe switching between the two mechanisms. The bottom panel of Fig. 7 shows the path length histograms when the increasing and decreasing transitions are distinguished. The decreasing subtrajectories from the fixed-length sampling and the trajectories from the flexible-length sampling (which are all decreasing) show much closer agreement, and the increasing transition shows a very different distribution.

We can also replace other common analyses with versions based on `Ensemble.split`. For example, consider the lifetime in a given state, which is defined by the time from when a trajectory first enters the state (having previously been in another state) until it enters another state. We refer to the desired state as *A* and the combination of all other states as *B*.

To think of this in terms of path ensembles, we describe it in two stages. First, we need to find the path ensemble which goes from another state, enters the desired state, and then enters another state. We denote this the “BAB” ensemble. The trajectories which are relevant to the lifetime calculation are subtrajectories of trajectories in the BAB ensemble. These go from the first entrance in A to the first entrance in We’ll call this the “AB” ensemble. We obtain trajectories in the AB ensemble by first getting all the segments that satisfy the BAB ensemble, and then selecting the relevant subtrajectories. Defining the BAB ensemble as

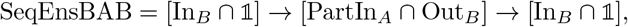

the corresponding OPS code is given by:

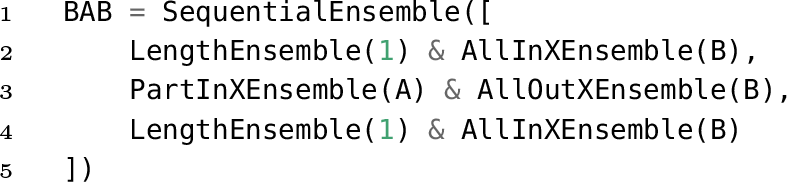

The AB ensemble is defined as

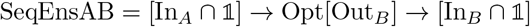

with the OPS code given by:

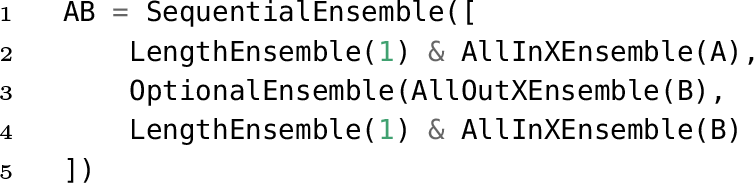

Both ensembles make use of the guideline on thinking about the halting criteria (from Sec. II G) by using Out_*B*_ as part of the middle subensemble. This guarantees that the middle subensemble stops, and the first frame afterward must be the first frame in state *B* (thus satisfying the final subsensemble). Additionally both of these ensembles use anchors requiring one frame in a state. Formally, the intersection with 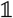 is not necessary, as we might as well have selected the last subtrajectory of frames in *B*/the first subtrajectory of frames in *A*, instead of the last frame in *B*/first frame in *A*.

To use these, we just run:

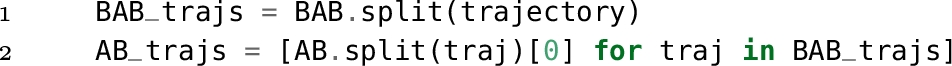

In Fig. 9, we visualize the trajectory segments associated with each of these ensembles for a sample trajectory from a toy model.

**FIG. 9.**
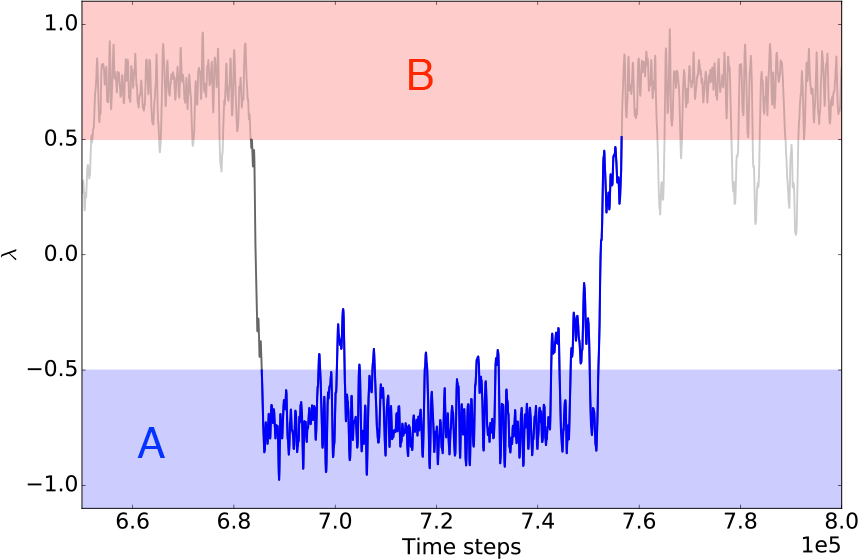
Trajectory segments for lifetimes. A trajectory for a toy model is shown with a light grey line. The red area represents state *B*, and the blue area represents state *A*. The segment that satisfies the BAB ensemble is shown in dark grey line, and the segment that satisfies the AB ensemble is shown in blue (and overlaps the BAB segment).

In the above example, the average time of the resulting trajectories gives the average lifetime in a state. In a two-state system, the reciprocal of the average lifetime is the rate. This could be modified to get the transition rate constants for a multiple state system by replacing the `AllInXEnsemble`(B) in the BAB sequential ensemble with an ensemble that would allow any state other than A, while the last one allows a specific state B. This would give the lifetime associated with the rate of *A* → *B*.

A similar procedure can be used to obtain the flux from a state through a given interface in TIS. In that case, we do two lifetime analyses: the lifetime outside the interface (where *A* everything outside the interface, and *B* is the state) and the lifetime inside the state (where *A* is the state, and *B* is everything outside the interface). The reciprocal of the sum of the average lifetimes from these gives us the flux [25].

Since this example involves two loops over the snapshots, it may not be as fast as code custom-designed to this purpose. (Although, in fact, the caching of OPS collective variables renders the vast majority of the total computational effort done in the first pass.) However, our primary intent here is to highlight how simple it is to prototype a trajectory analysis based on using the `Ensemble.split` method. It may be possible to write faster code, but it is hard to write code faster.

The specific implementations of flux and lifetime discussed here are included in the `SingleTrajectoryAnalysis` object.

### C. Custom PathMovers

OpenPathSampling also facilitates the creation of custom move schemes. In this section, we present an example of how that can be done, and compare the sampling behavior of this custom move scheme to the default move scheme.

The default RETIS move scheme selects a type of move at random (shooting, replica exchange, etc.) and then attempts one move of that type (shooting in a single ensemble, replica exchange fora specific pair, etc.). But perhaps a move which does all the replica exchanges in sequential order, then does shooting moves on all the ensembles, and then does the replica exchange in the reverse order, would be more efficient. This move will satisfy detailed balance — the question is whether it is more efficient.

Forthe most part, this simulation is set up as in the examples in Paper I[24]. The specific potential energy surface is given by

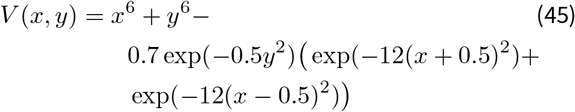

States are defined such that state *A* is *x* < −0.5 and state *B* is *x* > 0.5. Only the *A* → *B* transition is studied, with interface volumes from *x*_min_ = −∞ to *x*_max,*i*_ = {−0.4, −0.3, −0.2, −0.1}. The potential energy surface, with states and interface boundaries, is shown in Fig. 10.

**FIG. 10.**
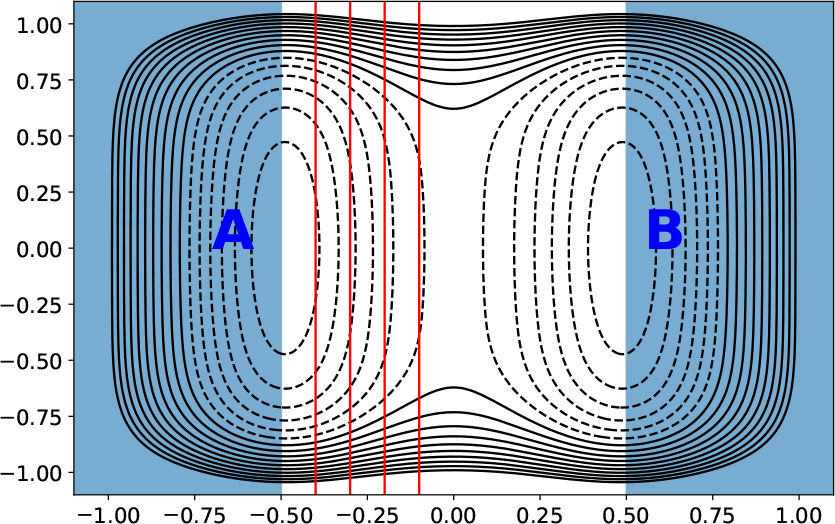
Modelsystem forthe custom move scheme example. Potential energy surface with states in blue and interface boundaries in red.

The main difference with the examples in Paper I[24]. is that here, we define a custom `MoveStrategy` object, which creates a custom sequential mover. The code for this mover and a move strategy to manage it is in Listing 2.

To use this strategy, we first create a default scheme, and then append the new strategy:

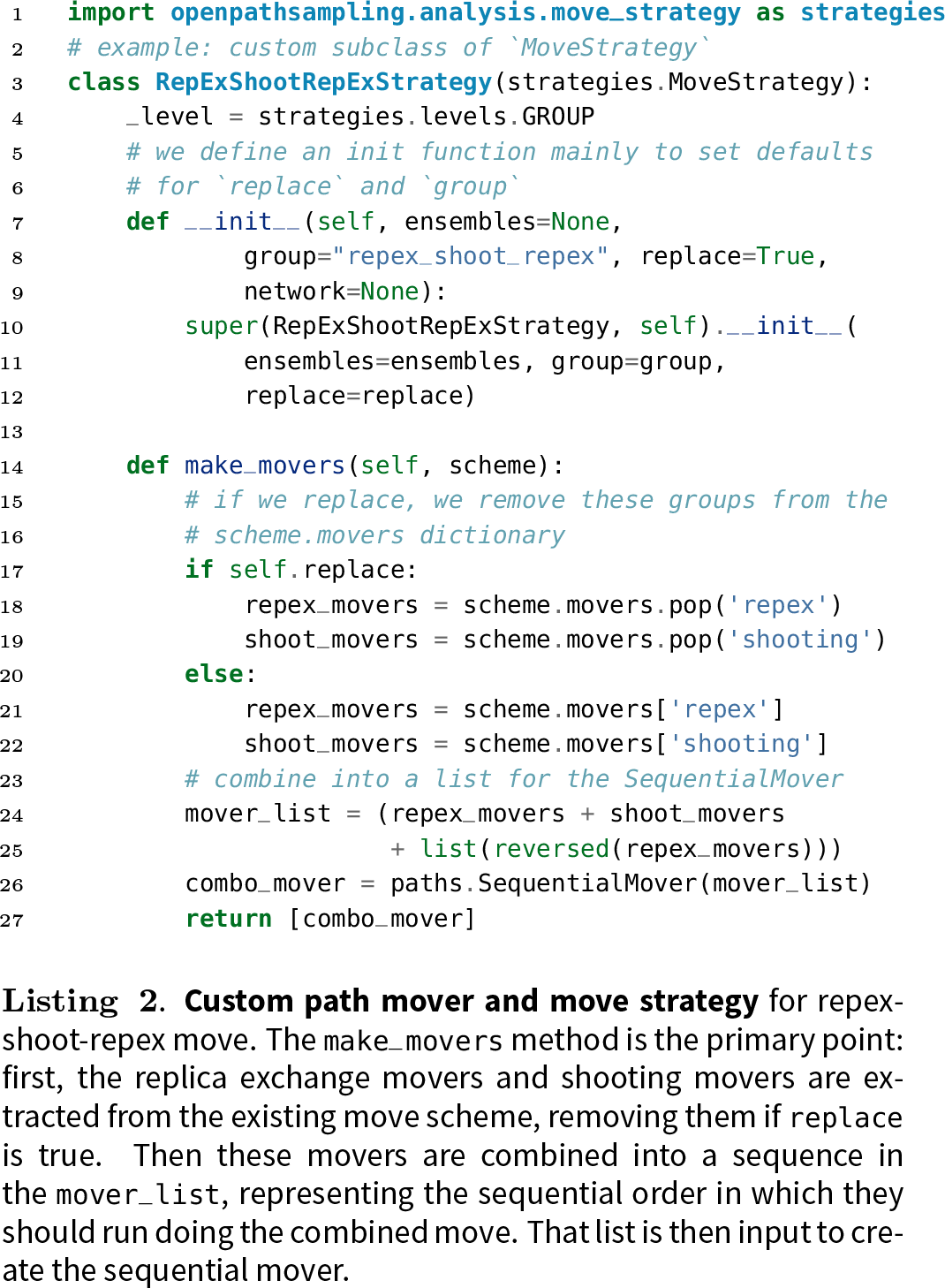

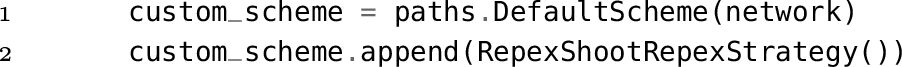

The visualization of this move scheme in is Fig. 11. There is still a random choice of type of move, but the types available are now minus move, path reversal, and the sequential mover, which is the only choice under the “`Repex_shoot_repexChooser`” in the illustration of the move decision tree. The relative probabilities for each move type are determined as ratios. By default, a new move type has the same (relative) probability as a shooting move, which is twice that of a replica exchange or path reversal, and five times that of a minus move. This means that when the repex-shoot-repex move replaces shooting and replica exchange in the move scheme, the ratio of per-ensemble shootingattemptsto path reversalsorto minus moves stays the same.

**FIG. 11.**
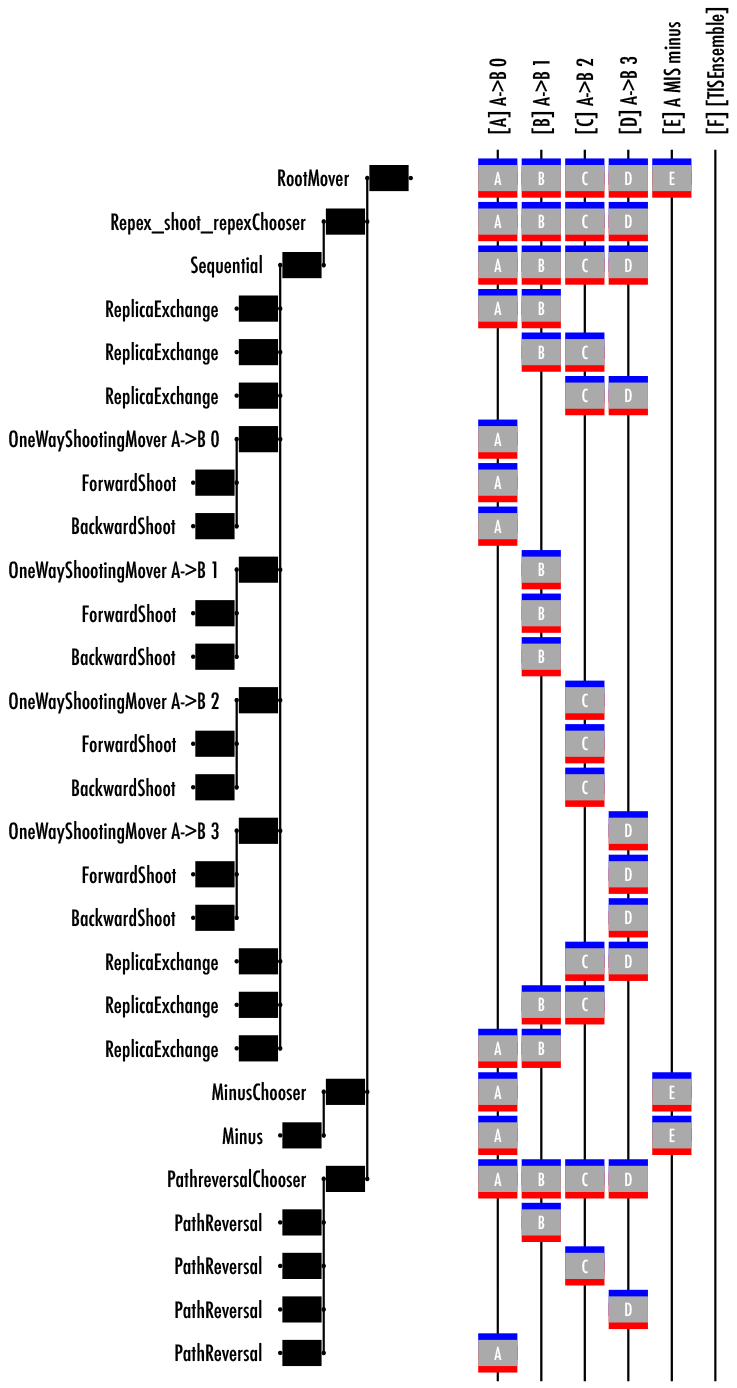
Move scheme including the repex-shoot-repex mover.

On the other hand, the number of total MC steps per ensemble shooting attempts will, of course, be very different between the custom and default schemes. To compare these fairly, we use the function `scheme.n_steps_for_trials`, which takes a mover and the number of desired attempts of that mover as arguments. We can ensure the two simulations do about the same amount of work by aiming for the same number of per-ensemble shooting moves. Forthe default scheme, this is

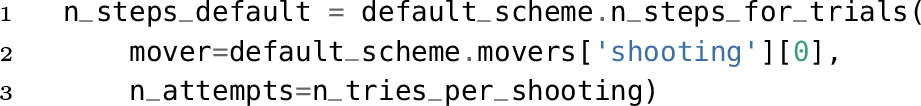

and for the custom scheme, it is

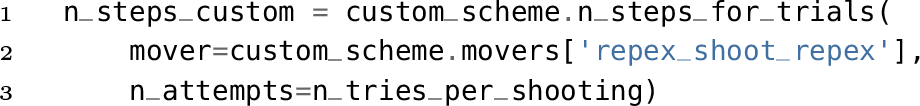

where `n_tries_per_shooting` is a number we have chosen (50000 in the example). We arbitrarily select the first shooting mover in the default scheme, since all have the same probability. We take the total probability of selecting the ‘`repex_shoot_repex`’ group, because there isonemoverin that group, and every time it occurs, it creates one shooting attempt foreach ensemble.

To ensure that this is a fair comparison, there are a few comparisons that should be made. First, we use the `scheme.move_summaryfunction` (described in PaperI[24])to show that we have the same number of path reversal and minus moves in each scheme. We can also use the move summary to see that we have the same number of shooting moves per ensemble, by comparing the per-ensemble shooting count obtained by adding ‘shooting’ as a second argument to `default_scheme.move_summary` with the number of `repex-shoot-repex` moves in the `custom_scheme`. Since both schemes are sampling the same number of shooting moves in the same ensembles, they should create roughly the same numberoftotalsnapshots. We can check this with len(`storage.snapshots`) for each storage.

To analyze these results, we consider replica round trip times and replica flow [34], concepts that monitorthe presence of bottlenecks during the replica exchange. Both concepts require defining ensembles as “top” and “bottom.” We put the minus ensemble as the bottom ensemble, and the outermost TIS ensemble as the top ensemble. A round trip can start from either the first entry to the “top” ensemble or the first entry to the “bottom” ensemble, and will use whichever the given replica visits first. If the replica visits the “top” ensemble first, the round trip duration is the number of Monte Carlo steps from the first entry into “top” until the replica returns to “top” after visiting “bottom,” with the case starting in “bottom” defined analogously.

Replica flow is defined by labelling each replica as either travelling “up” or “down,” depending on whether it more recently visited the “bottom” or“top” ensemble, respectively. For each ensemble *i*, the count of visits by “up” replicas is given by 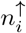, with the number of visits by “down” replicas given by 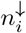. The flow for a given ensemble is defined as 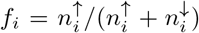. Formally, flow is 1 at the “bottom” ensemble and 0 at the “top” ensemble. The ideal flow is linear with the replica index. [34]

Round trip times and flow are both calculated as part of the `ReplicaNetwork` analysis tool. The code to analyze the default scheme is

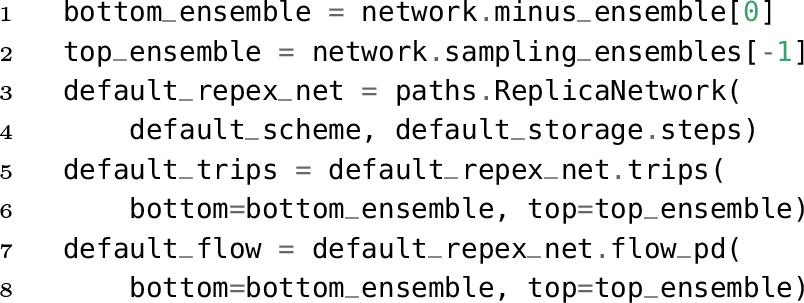

Then `default_trips`[‘round’] returns a list of the duration (in Monte Carlo steps) of each round trip that occurred. Analysis forthe custom scheme is analogous.

For this simple example, we find the custom move scheme does not yield significant improvement. The default scheme generates 345 round trips, whereas the custom scheme generates 311. There may be a small difference in the distribution of the round trip times (see Fig. 12). The distribution of round trip times is skewed toward slightly longer round trips forthe custom scheme. The replica flow, shown in Fig. 13, are very similar for both approaches. Overall, the default scheme is probably a slightly better choice.

**FIG. 12.**
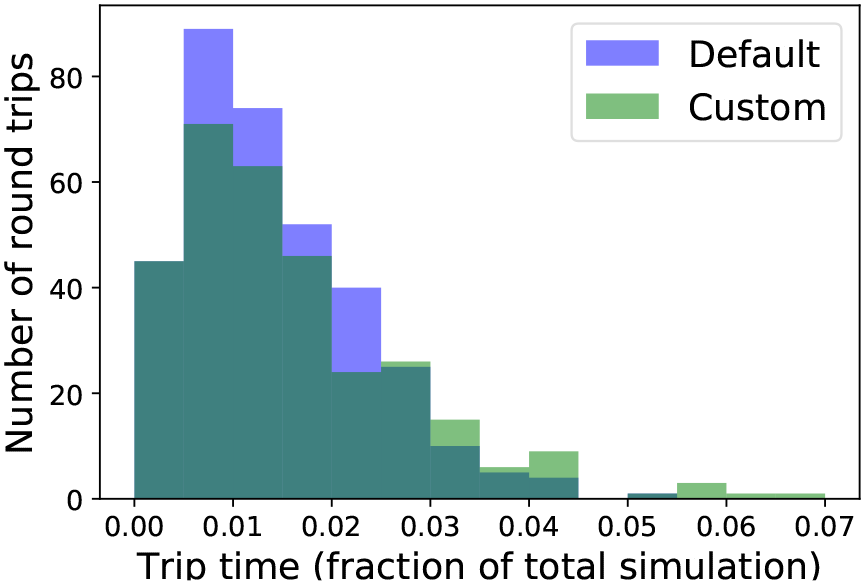
**Histogram of round trip times** for the default RETIS scheme and the custom scheme with the “repex-shoot-repex” move, with duration normalized to the total number of MC steps. Although both give about the same number of round trips, the distributions may differ somewhat.

**FIG. 13.**
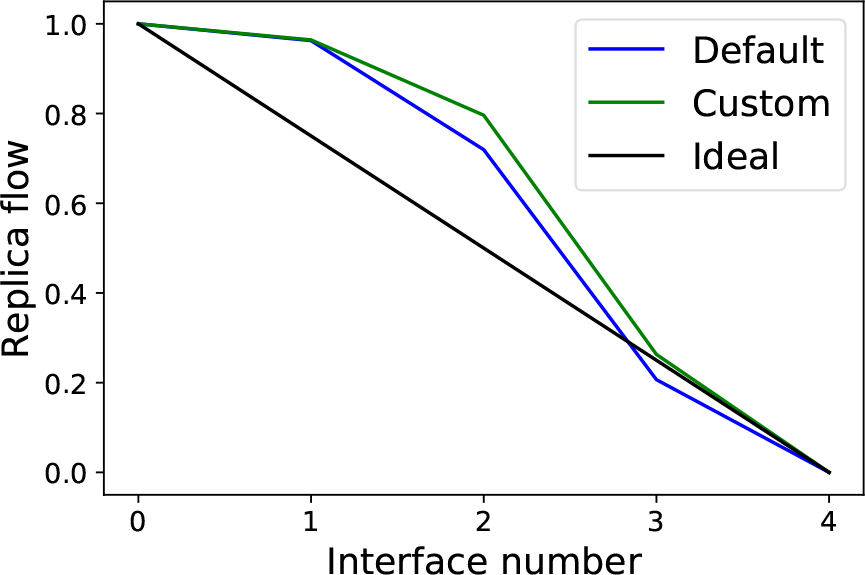
**Replica flow** for the default TIS scheme and the custom scheme with the “repex-shoot-repex” move. This example shows little difference, with the suggestion that the default scheme may be slightly better.

## V. CONCLUSION

In this paper we have described some advanced topics relevantto the OpenPathSamplingframework [24]. We have introduced a novel set-based approach to constructing path ensemble, along with a new notation suitable to this approach. This allows the application of set logic to path ensembles, and fits seamlessly with the way that OPS is written. Another advantage of this new notation is that it unifies the description ofthe monitorfunction of OPS with the path ensemble indicator function. Of particular importance herein is the sequential path ensemble, which is directly related to the way that OPS implements the path sampling monitoringand testing. Usingthis new notation it is remark-ablysimpleto create new path ensembles, and immediately implement these in OPS.

In addition, we provided insight in how one can customize the path sampling Monte Carlo movers within OPS in order to build non standard sampling schemes. These customizations are required if one wants to develop new path sampling schemes, or adapt existing ones.

In short, in this paper we have illustrated the power and flexibility of the OPS package. Users can now develop their own advanced sampling protocol entirely in OPS, and apply it to compute kinetic and thermodynamic observables.

In future work we will further elaborate on the foundation of the ensemble set-logic. Another direction is to parallelize the OPS code. While running multiple simulations in parallel is already possible, true parallelization requires the load balancing of multiple ensembles, where trajectories can have different and unpredictable path lengths, over the available computational resources.

## ACKNOWLEDGMENTS

DWHS and PGB acknowledge support from the European Union’s Horizon 2020 research and innovation program, under grant agreement No 676531 (project E-CAM). JDC acknowledges support from Cycle for Survival, NIH grant P30 CA008748, and NIH grant R01 GM121505. JDC, JHP, and DWHS gratefully acknowledge support from the Sloan Kettering Institute. FN acknowledges ERC consolidator grant 772230 “ScaleCell”, DFG NO 825/3-1, and SFB1114, project A04. project A04.

The authors are grateful for feedback from many people who helped beta-test the software, whose names are listed at http://openpathsampling.org/latest/acknowledgments.html. The authors are particularly grateful to Sander Roet (University of Amsterdam) for his feedback, and to Jocelyne Vreede (University of Amsterdam) forthe feedback obtained by using OPS as a teaching tool in courses on biomolecularsimulation.

